# Single Distal Mutation Enhances Activity of known PETases via Stabilisation of PET-binding

**DOI:** 10.1101/2024.09.11.612432

**Authors:** Xiaozhi Fu, Oriol Gracia I. Carmona, Gyorgy Abrusan, Xiang Jiao, Alexander Diaciuc, Mathias Gautel, Franca Fraternali, Aleksej Zelezniak

## Abstract

As a major source of plastic pollution, PET has attracted significant interest for biodegradation due to its potential in the circular economy. Finding effective enzymes still remains a challenge as screening methods are limited by either the low throughput or dependence on alternative non-PET substrates due to PET’s insolubility. Here, we report a highly active, stable and robust enzyme, Fast_2.9, identified while directly screening for PET-degrading activity in mesophilic conditions using droplet-based encapsulation of PET nanoparticles with the throughput above 1 kHz. We identified a distal S269T mutation that improves activity in the majority of all known PETases with up to 400 times over wildtype, and more than twice of known engineered PETases, as tested on untreated post-consumer plastics. Microsecond time scale molecular dynamics analyses indicate that this distant mutation possibly influences residues near the substrate-binding cleft via a common mechanism across PETases. Compared to the state-of-the-art FastPETase and LCC^ICCG^ enzymes, the engineered Fast_2.9 enzyme requires up to 8 and 42 times lower enzyme concentrations to reach the same enzymatic activity, ultimately requiring significantly less enzyme. As such our engineered enzyme degrades multiple post-consumer PET substrates, including polyester textiles, within as least as just 2 days with up to nearly 100% terephthalic acid conversion using as little as 0.72 mg_enzyme_/g_PET_ at 50℃. Our study presents a universal methodology for direct screening of insoluble substrates at ultra-high-throughput and highlights the techno-economic potential of Fast_2.9 for PET depolymerisation.

## Introduction

Polyethylene terephthalate (PET) is a widely used plastic in our daily lives, and it accounts for over 10% of the world’s solid waste with a major portion originating from polyester textiles ^1^. Micro and nano plastics could enter human bodies, and a recent study showed that those plastic particles in blood vessels might be linked to a higher risk of heart disease ^2^. Although it was considered non-biodegradable, the discovery of *Is*PETase ^3^ and cutinases ^4^ has shed light on its potential biodegradation processes that could potentially be reintegrated into a circular carbon economy via rapid enzymatic depolymerization of PET, with subsequent repolymerization or transformation into various valuable products ^5–7^. As a consequence, the scientific community rushed to search for new PET degrading enzymes ^8–12^ (>100 enzymes have been identified according to the Plastics-Active Enzymes Database - PAZy ^13^), and to engineer existing ones to enhance the activity, especially *Is*PETase through rational design ^10,14,15^ and directed evolution ^16^. For instance, ThermoPETase ^17^ was generated through rational design, FastPETase ^18^ and DuraPETase ^18,19^ were produced using computational methods, while HotPETase ^20^ and DepoPETase ^21^ were identified using directed evolution.

A traditional method for improving enzyme function is to use directed evolution by starting from a natural protein and iteratively making a small number of mutations until the enzyme acquires the desired properties ^22,23^. The combination of microfluidics and fluorescence cytometry has proven to be highly successful for this ^24–28^, especially for secreted enzymes ^29^ and substrates that cannot cross the cell membrane ^30^ with sufficient throughput. However, significant challenges remain when performing the directed evolution of PET-degrading enzymes. These include the extremely low water solubility of the PET substrate and the lack of mapping between PETase variants (i.e. genotypes) and absorbance or fluorescence (i.e. phenotypes) for easy detection. Few approaches have been developed for the directed evolution of PET degradation enzymes to overcome such obstacles. However, these methods are mostly of low throughput or use alternative substrates other than PET ^20,21^. In the case of HotPETase, solid PET discs were used in 96-well plates ^21^, and approximately 2000 samples were laboriously analysed using HPLC every two days. Qiao *et al ^16^* developed a microfluidic-based high throughput screening approach for PETase discovery but using substrates other than PET. As such, coupling fluorescence signals directly to enzymatic plastic polymer depolymerization in high-throughput still poses substantial challenges. For instance, Zurier et al ^31^ developed a microplate-based screening method for PETase mutants by adding Fe(II) in the PET hydrolysis solution and measuring the fluorescence of the 2-hydroxyterephthalic acid (hTA) with a spectrophotometer ^31^. hTA is the product from the coupling reaction of TPA with hydroxyl radicals, and it has excitation/emission wavelengths at 310/420 nm ^32,33^, which is out of the range of commonly used flow cytometry, thus limiting its measurements to spectrophotometry. On the other hand, to overcome the obstacles associated with linking genotype to phenotype, biosensors ^26,34^ are frequently employed. Notably, for PET degradation studies, transcription factor (TF)-based biosensors ^35,36^ have been developed to detect its monomer TPA, offering a promising tool for monitoring enzymatic activity and substrate interaction directly.

Here we developed a microfluidics platform to enable high-throughput screening of PETase activity directly using PET as a substrate by encapsulating PET nanoparticles into droplets. We engineered *the Escherichia coli* strain to efficiently secrete PETase and convert the resulting monomer yield into a measurable GFP signal using a genome-integrated TPA biosensor. We successfully applied our platform to screen a random mutagenesis library of *Is*PETase, resulting in >20% of sorted cells with increased activities, among which a variant with single mutation S269T (named as M2.9 in the following context) showed the highest increase of catalytic activity of 22-fold compared with *Is*PETase. We further discovered a universal activity enhancing effect of this mutation in various *Is*PETase variants (ThermoPETase, FastPETase, HotPETase and DepoPETase), especially for FastPETase, its S269 mutant (named as Fast_2.9 in the following context) exhibited 144% increasement in monomer yield using untreated postconsumer PET substrate at 50°C and pH 8. Microsecond time scale molecular dynamics analyses suggest that this distant mutation likely stabilises conformations accommodating the substrate, by influencing residues near the substrate-binding cleft via a common PETase mechanism. When benchmarking against non-pretreated postconsumer PET, Fast_2.9 showed up to 8 and 42 times lower of reverse Michaelis-Menten kinetic ^Inv^Km, compared to FastPETase and LCC^ICCG^ ^4^ respectively, while ^Inv^K_cat_ values remained similar. Using just 0.72 mg_enzyme_ ・ g -^1^ of enzymes towards non-pretreated postconsumer PET substrate in 15 ml bioreactor, Fast_2.9 achieved 97.4 ± 1.43% of depolymerisation and 95.9 ± 1.4% of TPA content by the end of 72 h, compared to 77.0 ± 4.74% depolymerisation and 55.4 ± 0.1% TPA content for LCC^ICCG^. Finally, we showed that Fast_2.9 is also capable of degrading polyester textiles with over 75% depolymerisation rate yielding up to 90% TPA monomer after 3 days.

## Results

### Engineering the strain for sensing TPA and secreting PETases

We started by engineering the *E.coli* BL21 (DE3) strain carrying a transcriptional factor (TF)-based TPA biosensor ^36^ (Supplementary Table 1) that would convert the PET depolymerisation catalytic efficiency into a fluorescence signal for subsequent screening (Fig. 1a). To warrant the stable expression of the biosensor ^37^, instead of using plasmids, we integrated the TPA biosensor into eight genomic loci using CRISPR-based multicopy chromosomal integration approach ^38^ (Fig. 1b). After three rounds of transformation, the biosensor was combinatorially integrated into four genomic locations resulting in six combinations ranging from a single copy (locus 2 and 4) up to four copies integrated into loci 2, 4, 5, 6 (loci 2456 strain) (Methods, Supplementary Table 2). To ensure that the biosensor is responsive to our defined dynamic range of TPA concentrations, all six strains were tested by submerging them to TPA concentrations of 0, 1 and 5 mM, with the intention to detect even minor substrate concentration differences (Fig. 1c). The loci 45 strain showed the most significant sensitivity capable of distinguishing 1 mM and 5 mM concentrations (Welch’s t-test, p-value < 2.2e-16).

**Figure 1.**
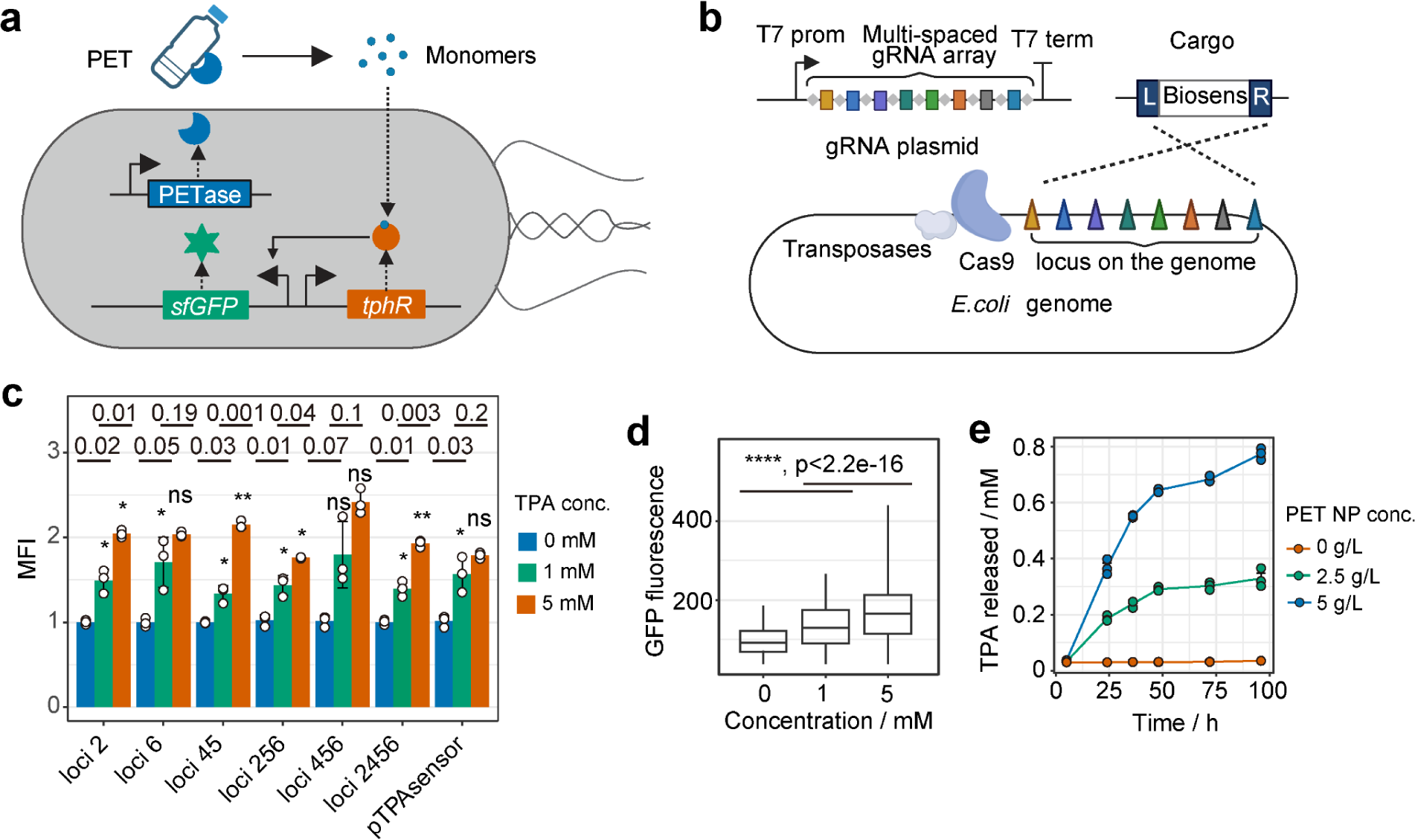
Co-expression of genomic based TPA biosensor and plasmid expressed *Is*PETase. **a,** Schematic representation of how the engineered strain converts PETase activity into fluorescence intensity. PETase is secreted to degrade PET outside the cell, and the resulting TPA monomer is taken up by the cell, inducing sfGFP expression downstream of the biosensor. **b,** Transposon-based CRISPR method for integrating the TPA biosensor into multiple genomic loci in *E. coli* BL21 (DE3). **c,** Median fluorescence intensity (MFI) ratio of different genomic-based TPA biosensor strains in OnEx System 2 medium supplemented with 1 or 5 mM Na2TPA, compared to the same strain cultivated in 0 mM Na_2_TPA. A strain expressing the TPA biosensor from a p15a ori plasmid (pTPAsensor) served as a control. MFI was calculated based on the fluorescence intensity of 5000 single cells measured by GUAVA EasyCite flow cytometry (n=3). Data are presented as mean ± SD. Statistical comparisons between samples at 1 and 5 mM TPA and the 0 mM TPA control were performed using a two-sided t-test with Welch’s correction. P values are indicated above each column, with NS, *, and ** denoting not significant, P < 0.05, and P < 0.01, respectively. **d,** Boxplot of GFP fluorescence distribution in the G-TPAsensor strain after 48 hours of cultivation in OnEx System 2 medium supplemented with 0, 1, or 5 mM Na_2_TPA, with 5000 events per group analysed by SONY SH800 cell sorter. The boxplot displays the median (line), interquartile range (IQR, box), and whiskers (rest of the data distribution). Statistical comparisons between different TPA concentrations were performed using a two-sided t-test, with P values <2.2e-16 indicated by ****. **e,** TPA yield of the G-TPAsensor strain when cultivating in the OnEx system 2 medium supplemented with 0, 2.5 and 5 g/L amorphous PET-NP from 0 to 96 h. Each group has triplicates. Each sample has triplicates and all data are shown as mean ± SD. Fig.1a and 1b were made using BioRender.com.

To corroborate whether the *Is*PETase enzyme was secreted and active in those strains, we transformed each strain with the plasmid-based T7-inducible *Is*PETase containing a N-terminal secretion signal peptide (Supplementary Table 3). In the strains where either locus 2 or 6 was targeted, no catalytic activity towards either PET or ρ-nitrophenyl acetate (a substrate for esterase activity) was ^39^ observed in the cell lysate or supernatant (Supplementary Fig. 1). This effect was independent of the use of expression mediums (we tested IPTG-supplemented induction and autoinduction mediums) (Supplementary Fig. 2). With both the TPA biosensor and secreted *Is*PETase functioning well, we continued with the loci 45 strain harbouring the pET21b-*Is*PETase plasmid, which is referred to as G-TPAsensor strain in the following context, unless otherwise specified.

We found that the fluorescence shift of the cell populations of G-TPAsensor in 0, 1 and 5 mM TPA remains significant after 48 hours cultivation (Fig. 1d), which confirms the biosensor’s functionality in long-term cultivation with plastics substrate and makes it suitable for identifying highly active mutants. Next, we cultivated the G-TPAsensor cells in the presence of 0, 2.5 and g/L PET nanoparticles (PET-NP) (made from amorphous PET, Methods) supplemented medium. The accumulations of monomers in 0, 2.5 and 5 g/L PET-NP exhibited linear growth in the first 48 hours and then increased slowly. 291 ± 2.4 μM and 645 ± 3.97 μM TPA were produced after 48 h (2.5 and 5 g/L PET, respectively), furthermore validating the functionality of the engineered strain (Fig. 1e).

### Accelerating PETase evolution in droplets

In order to cultivate the G-TPAsensor cells with PET substrate together within droplets, the PET substrate must be made into water-suspended nanoparticles. We first investigated the impact of nanoparticle preparation on PET substrate crystallinity ^40^, crucial for catalytic rates ^41^, by employing two methods in the Non-Solvent Induced Phase Separation (NIPS) process using hexafluoroisopropanol (HFIP) ^42^ and trifluoroacetic acid (TFA) ^43^ as solvents (Methods). We found that the PET nanoparticles obtained with TFA (PET-NP-TFA) method showed over double increased crystallinity as measured by differential scanning calorimetry (from 3.8% to 10.3%), while the PET-NP prepared with HFIP (PET-NP-HFIP) has had only 10% crystallinity increase (from 3.8% to 4.2%) comparing to the starting material (GoodFellow PET film, Methods). FT-IR analysis ^44^ also confirmed that the PET-NP-TFA has a higher crystalline conformation ratio with 52.5% increase of A1340/A1372 ratio and 20.0% decrease of A1475/A1457 ratio (1457 and 1373 cm^−1^ have been associated with the *gauche* conformation of PET, while 1475 and 1340 cm^−1^ increase in intensity during crystallisation due to *trans*) (Supplementary Fig. 3a and Supplementary Table 4). Therefore, we continued with the PET-NP-HFIP by encapsulating nanoparticles into water-in-oil-droplets (Methods).

We then encapsulated PET nanoparticles (diameter <500 nm, Supplementary Fig. 4) in water-in-oil droplets as a substrate together with *Is*PETase library to enable subsequent high-throughput PETase activity screening (Fig. 2a). Single cells from *Is*PETase random mutagenesis library (∼2 x 10^6^ clones) were encapsulated with PET NP within water-in-oil emulsions^46^ (Fig. 2b top). After 48 hours of droplets cultivation off chip, we observed cells grown together with PET-NP substrate exhibiting green fluorescence compared to negative control (Fig. 2b). We then de-emulsified the droplets and subsequently sorted cells using a fluorescent cell sorter. Initially, 77.1% of the cells were selected based on size and complexity criteria to exclude oversized or abnormal cells (Fig. 2c). Then, from this population, we isolated 192 cells with the highest green fluorescent protein (sfGFP) expression (Fig. 2d). The top-performing cells were then sorted onto agar plates in a 96-well format.

**Figure 2.**
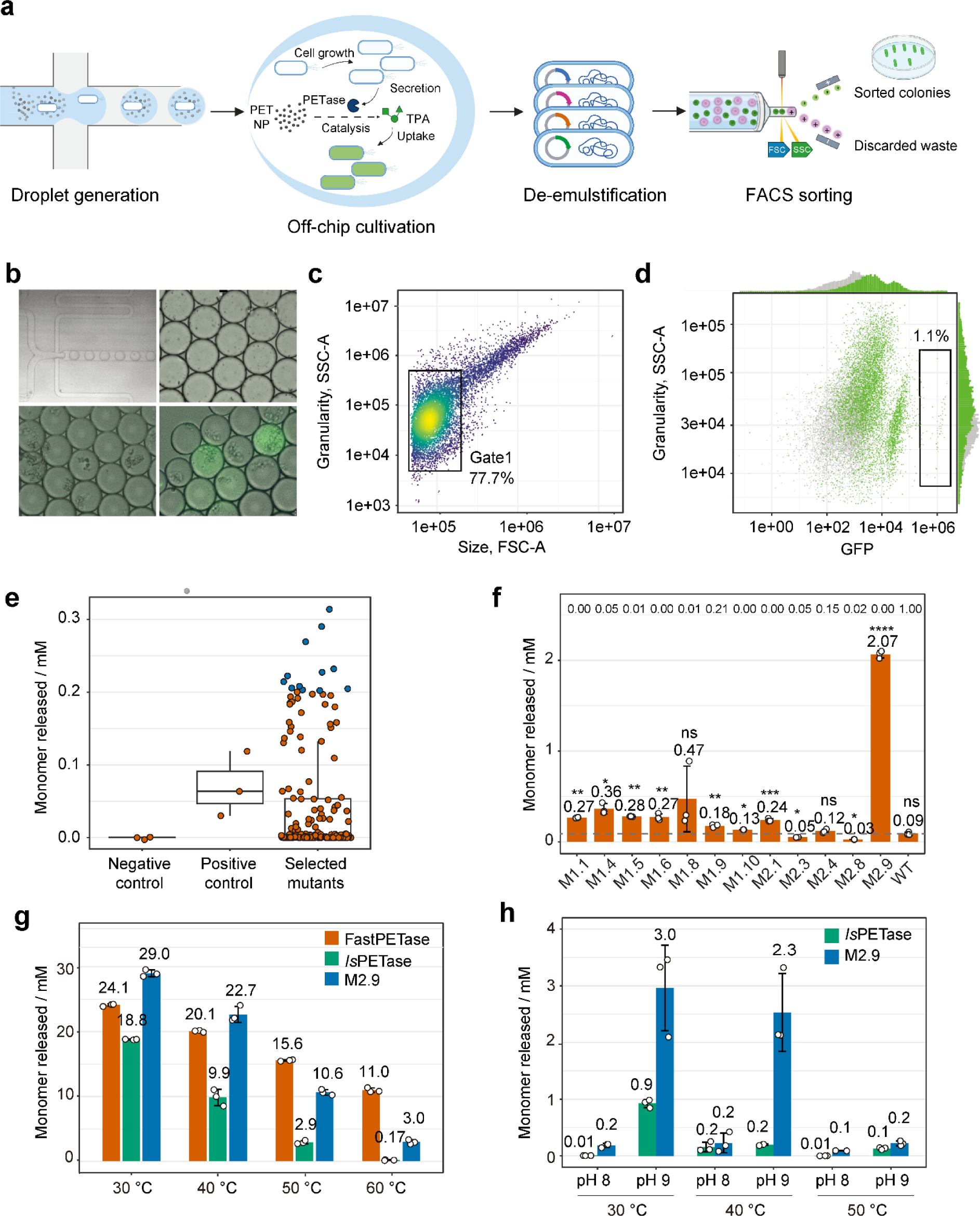
Droplet-based high throughput sorting platform for PET degrading enzymes. **a,** Single cells from a library of mutagenized PET-degrading enzymes were encapsulated in water-in-oil emulsions containing PET nanoparticle-supplemented medium. Following encapsulation, the cells were cultured off-chip within the droplets for 48 hours. After cultivation, the droplets were de-emulsified, and the cells were released for FACS sorting. **b,** Images showing droplets encapsulating both single cells and PET nanoparticles (PET-NP) generated on a co-flow droplet generator (top left), 60 μm diameter droplets after generation (top right), droplets from the negative control group (G-TPAsensor strain harbouring the pET21b plasmid) after 48 hours of cultivation in the fluorescent field (bottom left), and droplets from the library group after 48 hours of cultivation in the fluorescent field (bottom right). Images were taken twice, with the representative images shown here. **c,** The main population of 77.7% of cells was gated by FSC_A and SSC_A (Gate 1). **d,** The top 1.1% of fluorescent cells from Gate 1 were sorted. **e,** Boxplot of supernatant assay results from 167 selected mutants that grew on the sorting plate, alongside the positive control (G-TPAsensor strain harbouring the pET21b-*Is*PETaseWT plasmid) and the negative control (G-TPAsensor strain harbouring the pET21b plasmid). Each sample was run in triplicate, with jittered dots representing the mean value of each sample. The boxplot includes the median line, the interquartile range (IQR, box), and the whiskers denoting the rest of the data distribution. The top 12 samples, marked in blue, were selected for further purified enzyme verification, except for one that failed during purification. **f,** Barplot of purified enzyme assay results of the 12 samples selected from supernatant assay. Each sample was run in triplicate, and data are shown as mean ± SD. Statistical comparisons with the wild-type *Is*PETase (WT) were performed using a two-sided t-test with Welch’s correction. P-values are indicated above each column, with NS, *, **, ***, and **** representing not significant, P < 0.05, P < 0.01, P < 0.001, and P < 0.0001, respectively. **g,** Catalytic efficiency comparison of 200 nM *Is*PETase, *Is*PETase_S269T (M2.9) and FastPETase on 19.2 g/L PET-NP at 30, 40, 50 and 60 ℃ conditions. Each sample has triplicates and all data are shown as mean ± SD. Mean values of released monomers for each sample are indicated above each column. **h,** Catalytic efficiency of 200 nM *Is*PETase and *Is*PETase_S269T (M2.9) on 6 mM-diameter postconsumer transparent PET product (yoghourt container lid) under different temperatures (30, 40 and 50 ℃) and in different buffer conditions (100 mM KH_2_PO_4_ • NAOH pH 8.0 and 50 mM glycine • NaOH pH 9.0). Each sample was run in triplicate, and data are shown as mean ± SD. Mean values of released monomers for each sample are indicated above each column. Figure 2a was created using BioRender.com.

In total 192 variants were sorted out onto the LB agar and, of which 167 (86.9%) grew successfully on the sorting plate. Of these, 34 (20.3%) displayed greater catalytic efficiency than wildtype (Fig. 2e). We selected the top 12 of these 34 variants for sequencing (Supplementary Table 2) and further analysis. These 12 variants were purified and retested to confirm that their increased activity was due to higher catalytic efficiency and not merely due to elevated expression or secretion. Out of these, 6 variants showed significantly enhanced catalytic activities, with variant M2.9 (S269T) displaying the highest activity resulting in 22 fold increase over WT *Is*PETase variant (Fig. 2f). Variants M2.3 and M2.8, exhibited enhanced activity compared to the wild type in the supernatant assay. Yet, when purified, their activities were below that of the *Is*PETase wild type, likely due to the expression efficiency changes caused by the mutations ^35,36,47–49^.

To evaluate the catalytic efficiency of M2.9 we benchmarked it against FastPETase ^18^, which is known for its high efficiency among all IsPETase variants. 200 nM of FastPETase, *Is*PETase and M2.9 were mixed with 19.2 g/L PET nanoparticles at 30, 40, 50 and 60 ℃ for 48 h (Fig. 2g). M2.9 showed the highest activity at 30 and 40°C, producing 29.0 ± 0.526 mM and 22.7 ± 1.23 mM monomers, respectively. This represented increases of 20% and 13% over FastPETase, and 54% and 130% over *Is*PETase. A decrease in catalytic efficiency from 30 to 60°C was noted for all enzymes, with *Is*PETase and M2.9 displaying faster declines at 50 and 60°C compared to FastPETase, indicating that M2.9 is better suited for mesophilic conditions. Additionally, we tested M2.9’s activity on PET film from non-pretreated lids of postconsumer yoghurt containers (Supplementary Table 4). After incubating a 6 mm-diameter PET disc with 200 nM of the enzymes at 30°C for 48 hours, M2.9 produced 2.96 ± 0.753 mM of monomers, a 219% increase over the wild-type *Is*PETase (Fig. 2h). A more severe surface abrasion was also observed under optical and scanning electron microscopy analyses (Supplementary Fig. 5).

### Molecular dynamics and structural analysis of PETases

Residue 269 is located on the α7 helix of the surface of the protein ∼28 Å away opposingly from the active site consisting of residues Ser160, Asp206, and His237 ^50^ (Fig. 3a). Although the substitution of Serine for Threonine at this distal site is not expected to significantly impact the protein’s stability due to their similar properties, nonetheless, there are cases in other enzymes where the distant mutation from Serine to Threonine enhanced enzymatic activity, and also PETase^S242T^ that exhibited up to 2.5 increased activity compared with WT *Is*PETase^51,52^.

**Figure 3.**
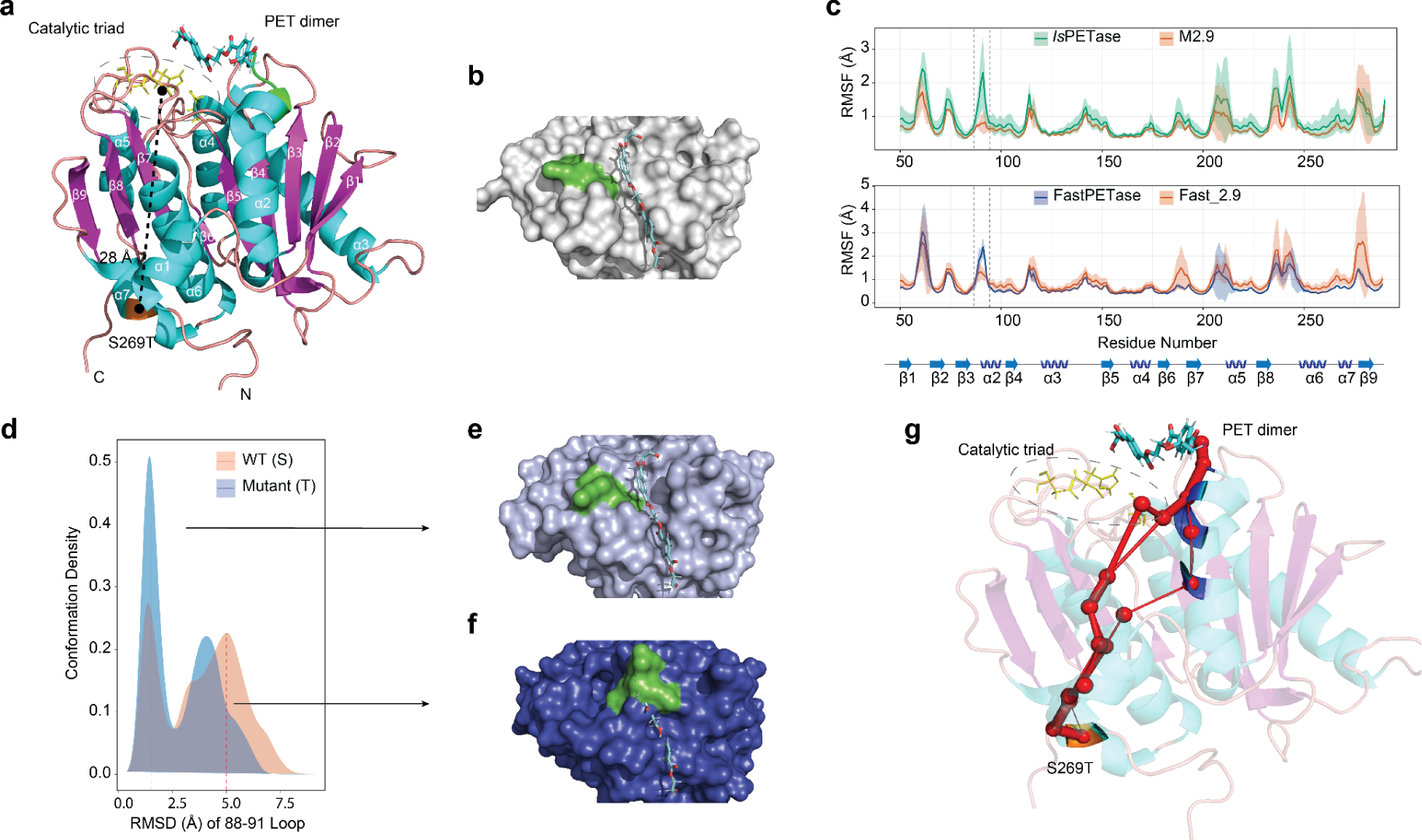
Molecular dynamics and structural analysis. **a,** Structure of *Is*PETase (cartoon structure) with a docked PET dimer (light blue), the catalytic triad (S160, D206 and H237) is highlighted in yellow and residue S269 is highlighted in orange. α-Helices are shown in cyan, β-sheets in magenta and loops in pink. **b,** Surface representation of *Is*PETase with the docked PET dimer in light blue. The loop region (residues 88 to 91) involved in the active cleft formation is highlighted in green. **c,** Average residue weighted Root Mean Square Fluctuation (RMSF) of the backbone atoms for the five replicates of PETase (green for *Is*PETase and blue for FastPETase), and the S269T variant (orange) for *Is*PETase (top) and FastPETase (bottom) simulations. The flexible loop region involved in the active cleft formation (residues 88 to 91) is highlighted in green. The S269T variants displayed significantly lower fluctuations than the wildtype versions (adj. p-value = 0.0264 and 0.0161 respectively). **d,** Probability density distribution of the RMSD of residues 88 to 91 relative to the crystallographic conformation (PDB code 7SH6 ^18^) for FastPETase simulation (orange) and the S269T variant (blue). **e,** Surface representation of a representative structure (estimated from 50 randomly sampled snapshots) of the FastPETase frames displaying low RMSD (RMSD ≃ 1.5 Å) for the loop region formed by residues 88 to 91 (green). The conformation of the PET dimer in the docked crystal structure is shown in light blue. **f,** Surface representation of a representative structure (estimated from 50 randomly sampled snapshots) of the FastPETase frames displaying high RMSD (RMSD ≃ 5 Å) for the loop region formed by residues 88 to 91 (green). The conformation of the PET dimer in the docked crystal structure is shown in dark blue. **g,** Residue correlation paths detected by WISP mapped onto the crystallographic structure of *Is*PETase. The catalytic triad is highlighted in yellow, residue S269 in orange and the docked PET dimer in light blue. Red spheres indicate the C-α of the residues involved in transferring the signal along the top 10 correlation paths estimated by WISP. The node edges indicate the detected paths, with width proportional to the frequency of the connection appearing in the top 10 paths.

To investigate the molecular effects of S269T mutation, we performed exhaustive molecular dynamics (MD) simulations, running 1 μs per replicate (n=5 per enzyme variant) allowing sufficient time for the mutation’s effects to propagate through the system (Methods). To locate the regions whose dynamics were affected by the mutation, we started by examining per residue weighted Root Mean Square Fluctuation (RMSF) profiles of the backbone atoms. As expected, most of the RMSF profile overlapped between the wild type *Is*PETase and S269T mutant simulations, with the exception of the following regions: the loops formed by residues 88 to 91 (unpaired t-test, BH adj. p-value = 0.0264), residue 112 to 115 (unpaired t-test, BH adj. p-value = 0.0287) and residue 238 to 241 (unpaired t-test, BH adj. p-value=0.0102). The region formed by residues 88 to 91 is both close to the active site and is involved in the formation of the active cleft where the substrate binds and has been shown to be a key difference between *Is*PETase and other members of the α/β-fold hydrolases ^53^ (Fig. 3b).

Given the high sequence similarity between *Is*PETase and FastPETase, particularly the identical residues from 88 to 91 in both enzymes, we hypothesised that the S269T mutation in FastPETase might act through the same mechanism and produce similar effects as observed in *Is*PETase. To investigate this, we performed molecular dynamics (MD) simulations for both FastPETase and its mutant Fast_2.9, each lasting 1 µs with five replicates. The simulations for both enzymes were prepared and analysed using the same methodology applied to the *Is*PETase simulations, with the sole difference being their optimal temperature (Methods). The per residue weighted RMSF profiles of FastPETase and Fast_S269T displayed the same stabilising behaviour as the one observed for *Is*PETase (unpaired t-test, BH adj. p-value=0.0161) (Fig. 3c).

To corroborate if the lower fluctuations observed in the region of residues 88 to 91 for the S269T mutants are due to a stabilisation of the crystallographic orientation or due to a new conformation, the Root Mean Square Deviation (RMSD) of the backbone atoms of residues 88 to 91 was analysed. To properly capture different conformations with respect to the crystal structure and not merely internal orientations of the loop, we aligned all simulation frames to the crystal structure using the C-alpha carbons of all residues except for 88 to 91. Following this alignment, we computed the RMSD for the backbone of residues 88 to 91 without additional realignment. We then generated a probability density distribution by combining data from all replicates. For both *Is*PETase and FastPETase, the S269T mutants showed a higher preference for conformations resembling the crystallographic structure, suggesting that S269T mutant acts by stabilising this conformation (Fig. 3d).

We then aligned the previously reported structure of *Is*PETase with a hydroxyethyl-capped PET dimer docked ^53,54^ with a representative structure of the loop conformations displaying low deviations with the crystallographic pose (Fig. 3d, RMSD ≃ 1.5 Å) and a representative structure of the loop conformations with a high deviations (Fig. 3d, RMSD ≃ 5 Å). The active cleft of the structure with lower loop RMSD preserved the active site cavity allowing the PET to fit without any major clashes (Fig. 3e) while the structures with high RMSD values blocked the active cleft cavity impeding the substrate from binding (Fig. 3f). Based on these findings, we hypothesise that the increased activity observed in both *Is*PETase and FastPETase resulting from the S269T mutation is attributed to the stabilisation of conformations that better accommodate the polymer substrate.

Since S269 is structurally close to C273 (∼4.8 Å), which forms a disulfide bond (DS2) with C289 (Supplementary Fig. 6a), we also investigated the DS2 and its possible interaction with S269. Although we did not find any direct evidence indicating the correlation between S269 and DS2, instead we found that the DS2 could possibly be partially reduced (Supplementary Figs. 6b, 7, 8), and the absence of DS2 did not significantly affect the activity of *Is*PETase but caused almost complete loss of activity in FastPETase (Supplementary Fig. 6c). We infer that in the WT *Is*PETase, the DS2 is likely to play a regulatory rather than structural role and further studies will be needed to clarify functions (See also in Supplementary note).

To get insights how the effect from S269T mutation may be traversing to ∼28 Å away active site, we analysed the shortest residue C-alpha fluctuation correlation paths connecting mutation to the residues of the active cleft using Weighted Implementation of Suboptimal Paths (WISP) method ^55^. As suggested by WISP analysis (Methods), the effect of mutation seems to propagate from the α7 helix to the α1 helix, then to the α2 helix and finally to the active cleft most commonly involving residues THR88, ALA89, SER93, ILE94 and TRP97 (blue residues in Fig. 3g).

### S269T enhances enzymatic activity across known PETases

As suggested by molecular dynamics analysis, we introduced the S269T mutation to FastPETase, an enzyme that is known to be active at higher temperatures^18^, to assess its effect on PETase activity. The mutated Fast_S269T variant showed enhanced catalytic efficiency across a pH range of 6-10, notably doubling its performance at pH 8 compared to FastPETase (Supplementary Fig. 9). Analogously, we applied the same S269T mutation to other *Is*PETase variants, i.e., ThermoPETase, HotPETase and DepoPETase, to see if this mutation exhibited universal effects (Fig. 4a). Similarly as with FastPETase, catalytic enhancements were observed in ThermoPETase across 30, 40 and 50 ℃ in pH 8 and 9 conditions. For HotPETase and DepoPETase, significant increases were only observed for its S269T variant at 50 °C pH 9 conditions (Welch’s t-test, p-value=0.00324 for HotPETase and p-value=0.0295 for DepoPETase), which possibly could be attributed to the structural differences of HotPETase and DepoPETase, which contain 21^20^ and 7^21^ mutations respectively, compared to *Is*PETase. The highest effects are observed in FastPETase overall showing over 430-fold increase of activity compared to wild type *Is*PETase performance at 50 ℃ and we named this enzyme Fast_2.9 (Fig. 4a).

**Figure 4.**
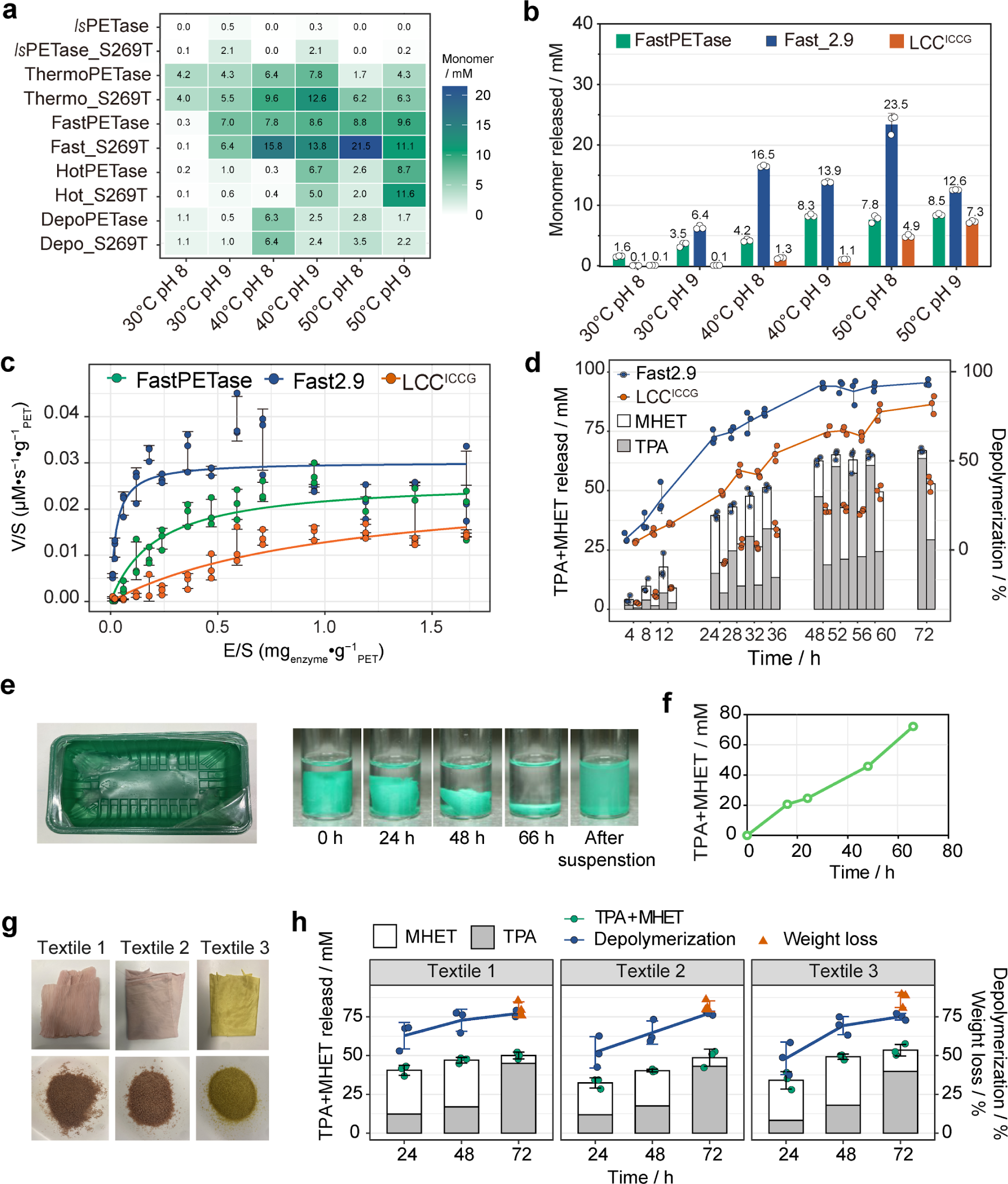
The S269T mutation enhances the catalytic performance of *Is*PETase variants. **a,** Heatmap showing the monomer release after incubating 200 nM *Is*PETase, ThermoPETase, FastPETase, HotPETase, DepoPETase, and their S269T variants with 6 mm-diameter postconsumer PET discs under various pH and temperature conditions for 48 hours. The experiment was performed in triplicate, and the mean values of released monomers are displayed in the heatmap. **b,** Total monomer release after incubating 200 nM enzymes with a 6 mm-diameter yoghourt container lid disc in pH 8 or pH 9 buffer at 30, 40, 50, and 60°C for 48 hours. All samples were run in triplicate, and data are shown as mean ± SD. The mean values of released monomers for each sample are indicated above each column. **c,** Inverse Michaelis-Menten plots of Fast_2.9, FastPETase and LCC^ICCG^ at 50°C in pH 8 buffer. All samples were run in triplicates and all data are shown as mean ± SD. **d,** Monomer yield and depolymerization rate of 12 g/L postconsumer PET disc by 100 nM (equal to 0.24 mg enzyme/g PET) Fast_2.9 and LCC^ICCG^ in 15 mL 100 mM KH_2_PO_4_・ NaOH buffer (pH 8) in Pioreactors over 72 hours. Additional enzyme portions (0.24 mg enzyme/g PET) were supplemented at 24 and 48 hours. All samples were run in triplicate, and data are shown as mean ± SD. **e, f**, A 40.6 mg piece of non-pretreated green opaque PET plastic was completely degraded by 0.0288 mg Fast_2.9 within 66 hours, with 0.0096 mg (0.24 mg enzyme/g PET) Fast_2.9 added at 0, 24, and 48 hours. Images were taken twice, and the representative images are shown. **g,** Images of three kinds of polyester textiles (Textile 1: outer skirt of a pink polyester dress; Textile 2: inne skirt of a pink polyester dress; Textile 3: yellow polyester T-shirt; top: raw textiles; down: textiles after pretreatment). **h**, monomer yield, depolymerisation rate and weight loss when incubating 12 g/L pretreatment textiles with 100 nM Fast_2.9 (which is equal to 0.24 mg enzyme/g PET) in 0.6 mL 100 mM KH_2_PO_4_・ NaOH buffer (pH 8) in 1.5 mL tubes within 72 hours. Additional enzyme portions (0.24 mg enzyme/g PET) were supplemented at 24 and 48 hours. All samples were run in triplicate, and data are shown as mean ± SD.

To access Fast_2.9 performance, we included LCC^ICCG^, a highly efficient enzyme tested in industrial scales ^56^. Since FastPETase and Fast_2.9 loses activity at 60°C (data not shown), we compared monomer production after 48h at temperatures 30, 40 and 50°C and pH levels of 8 or 9, revealing that Fast_2.9 is most effective at milder temperatures of 50 °C and pH 8, where it released 2.01 and 2.79 times more monomer FastPETase and LCC^ICCG^, respectively (Fig. 4a). Given these results, Fast_2.9 shows promise as a candidate for large-scale applications, especially at temperatures around 50°C, aligning well with the thermal properties of polymer, as isothermal crystallisation could possibly occur when the temperature is above the glass transition temperature (Tg) of the PET material ^41^ (Tg is 67 °C for amorphous PET and 81 °C for crystalline PET ^57^).

Enzymes have varying activities at different substrate concentrations, yet most of the studies utilise different enzyme-to-substrate ratios complicating benchmarking of enzyme performances. For instance, a non-saturated concentration of 0.2 mg_enzyme_・g_PET_^-1^ and a saturated concentration of 1 mg_enzyme_・g_PET_^-1^ was used in a comparative study of LCC^ICCG^ with FastPETase and another two PET degrading enzymes ^56^, while 200 nM and 11.9 mg PET was used in 600 μl buffer (equal to 0.282 mg_enzyme_・g_PET_^-1^) to compare FastPETase with LCC^ICCG^ and other enzymes ^18^. To investigate the effects of degradation rates in detail, we compared Fast_2.9, FastPETase and LCC^ICCG^ kinetics using the inverse Michaelis–Menten approach ^14^ using a range of enzyme concentrations. For all experiments, we used a unprocessed post-consumer PET substrate at 12 g/L across all conditions (Methods), with enzyme concentrations ranging from 5 nM to 700 nM (equivalent to 0.01 to 1.66 mg_enzyme_・g_PET_^-1^), measured over the first 7 h at a constant temperature of 50 °C and a pH of 8.

As shown in Fig. 4c and Table 1, we observed significant differences among the three enzymes. While ^inv^V_max_ /S value of Fast_2.9 is 15.4% higher than both FastPETase and LCC^ICCG^, especially striking is ^Inv^Km differences, which is 8.5 and over 40 times lower than the ^Inv^Km values of FastPETase and LCC^ICCG^ respectively, indicating much less of enzyme for Fast_2.9 is needed to fully depolymerise PET. We did notice a slight decrease in the velocity when the enzyme concentrations were higher than 0.95 mg_enzyme_・g –^1^ for Fast_2.9 and 1.19 mg_enzyme_・g –^1^ for FastPETase, which has also been reported for *Is*PETase before ^58^. This concentration-dependent inhibition, likely due to surface crowding ^58–61^, is a common behaviour in the enzymatic degradation of solid substrates, as observed in cellulases and polyhydroxyalkanoate (PHA) depolymerases^60^. Such phenomenon is much less pronounced in enzymes with higher thermal stability likely due to higher rigidity ^59^, which explains the absence of such concentration-dependent inhibition for LCC^ICCG^ (Fig. 4c).

**Table 1.** Kinetic parameters of Fast_2.9, FastPETase and LCC^ICCG^ derived from the inverse Michaelis‒Menten experiments.

| Inverse MM kinetics | $^{inv}K_M$ (mg<sub>enzyme</sub> • g<sub>PET</sub><sup>-1</sup>) | $^{inv}V_{max}/S$ (μM • s<sup>-1</sup> • g<sub>PET</sub><sup>-1</sup>) |
| --- | --- | --- |
| Fast_2.9 | 0.025 | 0.030 |
| FastPETase | 0.213 | 0.026 |
| LCC<sup>ICCG</sup> | 1.049 | 0.026 |

Enzyme costs play a significant part in the economics of industrial enzymatic hydrolysis ^14,35,36,62^. The lower in ^inv^K_M_ of Fast_2.9 indicates a lower amount of enzyme is needed, thereby reducing the overall recycling process costs. We next applied the least saturated enzyme concentration of Fast_2.9 (0.24 mg_enzyme_・g_PET_^-1^) to degrade 7 types of untreated postconsumer PET products (Supplementary Table 4), including two coloured - black and green opaque PET (Supplementary Fig. 10). We observed that 4 mg of post consumer PET could be completely degraded in as little as 22h (Supplementary Fig. 10a). Using the same enzyme amount (0.24 mg_enzyme_・g_PET_^-1^ of enzyme was added at 0, 24, 48 h, which means in total 0.72 mg_enzyme_・g_PET_^-1^ of enzyme), analogously, we then compared Fast_2.9 to LCC^ICCG^ by scaling the reaction to 15 mL volume at 50 °C in pH 8 buffer (Fig. 4d). We found that Fast_2.9 exhibited a higher depolymerization rate compared with LCC^ICCG^ during the entire 72 h incubation period. The accumulated monomers of TPA and MHET of Fast_2.9 reactions reached its plateau at around 48 h, while for LCC^ICCG^ the total monomer concentrations kept increasing until the end of experiment without reaching complete depolymerisation even at the final 72 hours point. We also found that the TPA fraction of total monomers in Fast_2.9 reactions kept increasing during the last 24 h and reached 95.9 ± 1.4% by the end of 72 h, while the final TPA ratio in total monomers in LCC^ICCG^ reactions was 55.4 ± 0.1%.

Most transparent PET bottles are mechanically recycled in developed countries, but not coloured containers ^63^. Therefore we also evaluated how Fast_2.9 could degrade untreated coloured opaque postconsumer PET containers with just 0.72 mg_enzyme_・g –^1^ of enzyme usage. Under identical conditions, 97.6% of PET was degraded within 66 hours leaving only the green pigment in the reaction container (Fig. 4e, 4f, Supplementary Video) demonstrating Fast_2.9 suitability for TPA recovery from non-transparent PET waste.

Approximately 60% of the PET produced is used in the textile industry, while ∼30% is used in the PET bottle industry ^64^. And there exist no mature recycling system for textile waste globally, 87% of which ends in incineration or landfills ^65^. To explore the broader application of Fast_2.9, we pretreated 3 types of polyester textiles with extruding, quenching and grinding (Fig. 4g), and then incubated with Fast_2.9. By the end of 72 hours of incubation, the depolymerisation rate reached 77.2 ± 1.6%, 77.4 ± 1.1% and 75.1 ± 1.8% respectively, which is also close to the substrate weight loss of 79.7 ± 4.0%, 81.8 ± 2.8% and 85.9 ± 4.0%, as measured by lyophilization weight loss (Fig. 4h). And similar to PET bottle degradation in Fig. 4d, the TPA ratio in total monomers increased in the last 24 hours, which reached 89.9 ± 1.8%, 89.2 ± 9.5% and 74.4 ± 4.0% respectively (Fig. 4h).

## Discussion

In this study we presented a novel droplet-based ultra-high throughput screening approach that enables direct use of insoluble PET substrates for directed PETase evolution to surpass previous limitations of low throughput screening. We engineered the strain using a biosensor that is TPA specific^35^ and transformed PET into nanoparticles to allow directly to sense PET degradation without relying on alternative substrates that are not related to depolymerisation of PET^16^. Through screening a million-sized library at mesophilic 30°C temperature, we identified eight mutants that enhanced *Is*PETase activity, with the most notable increase, a 22-fold effect (Fig. 2f), observed with the single S269T mutation located on the α7 helix, that is distant from the active site (Fig. 3a). We introduced the S269T mutation into known PETases and observed a general trend of activation, most pronounced in FastPETase (Fig. 4a). The resulting enzyme, Fast_2.9, achieved a 430-fold increase in activity over the wild type at 50°C using untreated post-consumer substrate (Fig 4a).

To explore the underlying mechanisms, we conducted extensive molecular dynamics simulations as distal mutations might influence protein structure and dynamics through long-range effects, potentially leading to allosteric impacts on enzymatic catalysis ^66–69^. Molecular trajectory analysis revealed that the ∼28Å distance is mediated by interactions involving residues that connect to the β3-α2 loop, stabilising residues Y87, T88, and A89 (Fig. 3c, 3g). Notably, Austin et al^53^ found that the substrate binding cleft of *Is*PETase includes T88 and S238, while Guo and colleagues^70^ demonstrated that T88 and A89 are crucial for interacting with the PET polymer, such that mutations Y87H, Y87E, T88H, and A89H resulted in a significant loss of activity, underscoring the critical roles these residues play in enzyme function. Similar to *Is*PETase and M2.9, we found the same global change pattern for FastPETase and Fast_2.9 (Fig. 3c, 3d, 3e, 3f).

PET typically has a glass transition temperature (Tg) starting from 67°C for amorphous PET, meaning reactions at higher temperatures can lead to increase of PET crystallinity^71^ thereby preventing PETase catalysis. Despite that additives like plasticizers and exposure to water can lower its Tg ^72,73^ still it is important to consider these as PET may recrystallize if the environmental temperature exceeds its Tg ^74^. For instance, Thomsen and colleagues observed that the Tg of amorphous PET disks decreased from 75°C to 60°C after a 24-hour pre-soak in water at 65°C ^41^. In fact, Arnal et al tested the enzyme variant LCC^ICCG^, a variant known for its high efficiency in industrial applications, at a lower temperature of 68°C instead of 72°C, taking into account the thermal properties of PET ^56^. Our results show that Fast_2.9 has higher activities than FastPETase at 30-50 ℃ (Fig. 4a, 4b) and in different pH (Supplementary Fig. 9), also outperforming LCC^ICCG^ in temperature ranges 30-50 ℃ (Fig. 4b), and thus aligning well with the thermal properties of PET and making overall favourable process economy in mesophilic temperatures.

Inverse Michaelis-Menten kinetics analysis of the three enzymes with 12 g/L postconsumer PET plastics showed that Fast_2.9 has the lowest ^Inv^Km, which could be attributed to the substrate binding cleft stabilisation by S269T for better substrate binding (Fig. 3). Lower ^Inv^Km means lower enzyme usage, lowering costs of enzymatic catalysis, whereas the low affinity of LCC^ICCG^, as previously suggested, could potentially limit its wide industrial adoption^75^. As such, we showed that as little 0.72 mg_enzyme_・g ^-1^ of Fast_2.9 is needed for nearly 100% conversion into TPA of multiple postconsumer PET (both transparent and coloured opaque products) within just 48 hours, which is 28% less enzyme amount than the optimised 1 mg_enzyme_・g ^-1^ for LCC^ICCG^ ^56^ and 64% less than the recently reported 2 mg_enzyme_・g –^1^ for TurboPETase ^14^. Minimising enzyme usage in this process plays a critical role in enhancing the transformation of PET waste into high-value products, thereby contributing to the overall value enhancement of this recycling pathway. High conversion rate from raw PET to TPA rather than MHET is needed to achieve the circular biorecycling and is preferable to be done by a single enzyme. Here we showed the potential of Fast_2.9 acting as MHETase ^76^ and achieved 95.9% TPA content in the total monomers. Similar function has been observed in the variant *Is*PETase^S92K/D157E/R251A^ ^77^, however the exact content value was not reported. Furthermore, in all our experiments except for textile degradation, we used post-consumer PET from customer-used bottles and containers directly, without any pre-treatments such as heating, grinding ^18^, acid and base treatments ^78,79^, which underscores the application potential of Fast_2.9. In this study, secondhand polyester textiles were the only material subjected to pretreatment. These textiles typically exhibit higher crystallinity and contain more additives compared to food-grade bottles or containers. Initial attempts to directly degrade the raw textile using Fast_2.9 showed limited efficiency (data not shown). However, by incorporating a pretreatment step followed by enzymatic degradation at 50°C, we demonstrated the effectiveness of Fast_2.9 in recycling textile waste. This method achieved over 77% depolymerization and 74.4-90% end-to-end TPA production, highlighting its potential for sustainable textile waste management.

The resulting high activity, low substrate-enzyme ratio of Fast_2.9 and effectiveness at mild 30-50℃ temperatures makes it a suitable candidate for industrial PET depolymerisation. In PET enzymatic processing, a higher concentration of TPA is advantageous because an additional hydrolysis step is normally needed to completely convert MHET into TPA either by enzymatic ^80^ or chemical hydrolysis ^18^. TPA can be used as a fundamental building block for synthesising recycled virgin-like PET^4^ as well as for upscaled novel applications ^7,81^. As such, ∼96% TPA yield also justify advantageous industrial suitability of Fast_2.9 over state-of-the-art rivals (Fig. 4d). Furthermore, S269 and its derived mutants can be further enhanced through iterative cycles of our presented droplet-based enzyme evolution with substrate nano-particles. Overall, we foresee this technology as a general method that can be applied for developing a broad range of biocatalysts acting on water insoluble solid substrates including other environment polluting plastic polymers^82^. Accelerated enzyme evolution has once again transformed previously challenging enzymes, now, into potential drivers of the plastic circular economy.

## Methods

### Genomic integration of TPA biosensor in *E.coli*

The TPA biosensor module (Supplementary Table 1) was synthesised, cloned into pTwist Chlor Medium copy number plasmid by TWIST Bioscience (www.twistbioscience.com) and named pTPAsensor. Four plasmids (pDonor, pTnsABC, pQcascade-array1 and pCutamp) for multicopy chromosomal integration using CRISPR-associated transposases (MUCICAT) were purchased from Addgene (https://www.addgene.org). The pDonor-TPAsensor plasmid was constructed by inserting the TPA biosensor module into the pDonor plasmid. The TPA biosensor biomodule was amplified using the primer pair TPA sensor_pDonor_F and TPA sensor_pDonor_R (Supplementary Table 6) and gel-purified with GeneJET Gel Extraction Kit (Cat#K0691, Thermo Scientific). The purified PCR product was ligated into the linearized pDonor plasmid using Gibson assembly ^83^, following BamHI and XbaI digestion. The pDonor-TPAsensor plasmid was sequence-verified by Eurofins Genomics (https://eurofinsgenomics.eu/) using the primer pair pDonor_seq_F and pDonor_seq_R after colony PCR.

BL21(DE3) competent cells were co-transformed with pDonor-TPAsensor and pTnsABC plasmids. After a 1-hour recovery incubation, the transformed cells were plated on LB-agar plates containing 100 μg/mL ampicillin and 50 μg/mL kanamycin. The recombinant BL21(DE3) strain carrying pDonor-TPAsensor and pTnsABS was made chemically competent using the Hanahan Method ^84^ and then transformed with pQCascade-array1 plasmid. These cells were plated on LB-agar containing 100 μg/mL ampicillin, 50 μg/mL kanamycin, and 50 μg/mL streptomycin, and incubated at 37 °C for 16 hours, leading to colony growth.

These colonies were scraped, suspended in fresh LB medium, and a 100 μl aliquot was re-plated on LB-agar plates containing the triple antibiotic combination and 0.1 mM IPTG to induce transposition-related protein expression and biofilm formation. The plates were incubated at 37 °C for an additional 16 hours. After biofilm formation, the biofilms were scraped off, resuspended in LB medium, diluted, and plated on LB-agar with the triple antibiotic combination and 1 mM IPTG. These inducing plates were incubated overnight at 37°C.

To enhance clonal outgrowth and integration efficiency, colonies from the inducing plates were designated as Transfer 1. A subset of these colonies was streaked onto fresh LB-agar plates with the triple antibiotics and 1 mM IPTG, then incubated overnight at 37 °C, resulting in Transfer 2. Further transfers led to Transfer 3. Copy numbers were identified using colony PCR or qPCR with 8 primer pairs (cr1-cr8, Supplementary Table 6) targeting the genomic sites integrated by pQCascade-array1.

### Generation of random mutagenesis library

The DNA sequence of *Is*PETase (excluding signal peptide) was subjected to error prone PCR using varying concentrations of MnCl_2_ (0, 0.1, 0.2, 0.3 mM) to control the mutation rate, with 10 ng pET21-PETase as the template in a 50 μl PCR system. The primer pair PETase_RM_SP_F and PETase_RM_R (Supplementary Table 6) was used. The PCR products were purified by Genejet PCR purification kit (Cat#K0701, ThermoScientific) and subsequently used for the amplification of the pET21b-PETase plasmid via the RF cloning method ^85^. This involved mixing 100 ng of purified gene fragments with 20 ng of plasmid in a 50 μl PrimeStar HS polymerase PCR system (Cat#R010A, Takara) and following the PCR conditions: 98 ℃ for 30 s, then 15 cycles of 98 ℃ for 10 s and 68 ℃ for 6 min, followed by 72 ℃ for 10 min, with incubation at 16 ℃.

To digest the circular plasmid, 1 μl of DpnI (Cat#FD1704, ThermoScientific) was added to each 10 μl RF cloning PCR product, and the mixture was incubated at 37 ℃ for 30 minutes. The digested products were then electroporated into ElectroMAX™ DH5α-E Competent Cells (Cat#11319019, ThermoScientific) following the manufacturer’s protocol. All clones from each plate were pooled and cultured in LB medium, except for a few individual clones from each library, which were picked and verified by sequencing at Eurofins Genomics. Plasmids from the pooled culture were purified using the GeneJet Plasmid Miniprep kit (Cat#K0503, ThermoScientific) and electroporated into the G-TPAsensor strain competent cells..

### Site directed mutagenesis

DNA libraries targeting specific residue positions were constructed using the Q5 Site-Directed Mutagenesis Kit (Cat#E0552S, NEB) following the manufacturer’s protocol. To mutate S269 to T269, the following primer pairs were used: ThermoPETase_S269T_F / ThermoPETase_S269T_R for ThermoPETase, FastPETase_S269T_F / FastPETase_S269T_R for FastPETase, HotPETase_S269T_F / HotPETase_S269T_R for HotPETase, and DepoPETase_S269T_F / DepoPETase_S269T_R for DepoPETase. To mutat C273 to A273, the primer pair IsPETaseC273A_F / IsPETaseC273A_R was used for IsPETase, and FastPETase_C273A_F / FastPETase_C273A_R was used for FastPETase. To mutate C289 to A289, the primer pair IsPETaseC289A_F / IsPETaseC289A_R was used for IsPETase, and FastPETas_C289A_F / FastPETas_C289A_R was used for FastPETase. The primer sequences are provided in Supplementary Table 6. All plasmids were sequenced at Eurofins Genomics before proceeding to protein expression."

### Preparation of PET nanoparticles

For the HFIP method ^42^, 0.58 g GoodFellow amorphous PET film (Cat#ES30-FM-000145, GoodFellow) was dissolved in 35 mL hexafluoroisopropanol (HFIP) (Cat#AAA1274722, ThermoFisher) with magnetic stirring. PET solution (10 mL) was added dropwise into 500 mL ultrapure deionized water with vigorous stirring at room temperature, resulting in precipitation of PET-NPs. For the TPA method ^43^, one gram of PET particles was dissolved in 10 mL of concentrated trifluoroacetic acid solution (TFA) (90% v/v) at 50 °C and stirred until complete dissolution, which took approximately 2 h. After dissolution, the solution was left to stand overnight. To precipitate the micro/nanoparticles, 10 mL of a diluted aqueous solution of trifluoroacetic acid (20% v/v) was added to the initial mixture under vigorous stirring and kept stirred for 2 hours, followed by overnight storage. The suspension was centrifuged at 2500g for 1 h, and the supernatant was discharged. The pellet was resuspended in 100 mL of ultrapure deionized water with vigorous stirring at room temperature. For both methods, the PET NP suspension was placed in a fume hood at 55 °C with magnetic stirring to remove residual HFIP. A 1 mL aliquot of resuspened PET nanoparticles were lyophilized for concentration analysis and HFIP, DSC analysis. The PET NP suspension was always ultrasonicated for 10 min freshly before use.

### Droplet generation and manipulation

The cell suspension was diluted with Overnight Express OnEx system 2 (Cat#71300, Merck) medium supplemented with 2.5 g/L PET nanoparticles and 200 μg/ml ampicillin. Cells were diluted to an OD ∼0.1 to achieve an average 0.3 cell per droplet according to Poisson’s distribution ^45^, based on the assumption that 1 mL of this *E. coli* suspension at A600 nm = 1 would contain 5 × 10^8^ cells ^86^. Both the cell suspension and HFE 7500 (Cat#EU-FLUO-7500-10, Emulseo) with 0.5% FluoSurf Surfactant (Cat#EU-FSC-V10-2%-HFE7500, Emulseo) were pumped into a Co-Flow Droplet Generator (Cat#DG-CF-35-05, Droplet Genomics) with a flow rate of 10 μL/min by Flow EZ system (Fluigent).

The resulting 2 mL droplets were collected in a 15 mL sterilised falcon tube and kept at 30°C, 200 rpm for 2 days in a shaker. After 48 hours of cultivation at 30 ℃, 500 μL 1H,1H,2H,2H-perfluoro-1-octanol (Cat#370533-25G, Merck) was added into the emulsions and mixed gently until droplets were all de-emulsified. Following this, 1 mL sterilised PBS buffer (pH 7.0) was added and the mixture was settled for 5 min. The oil layer settled at the bottom, with the PET nanoparticles sedimented in the middle layer. The cell suspension, found in the upper layer, was collected for FACS analysis and sorting.

### Fluorescent microscopy

Droplets were suspended with HFE 7500 oil and deposited onto glass slides for microscopic analysis. Observations were conducted under both bright and fluorescent fields, using 20x and 40x objectives on a LEICA DM2000 microscope (Leica Microsystems CMS GmbH).

### Flow cytometry analysis by GUAVA easyCyte Flow Cytometer

For the functional analysis of plasmid-expressing and genomic-integrated biosensor strains, cells were analysed using a GUAVA easyCyte Flow Cytometer (Luminex). Cells were cultivated overnight in LB medium supplemented with 100 μg/mL ampicillin and 15 μg/mL chloramphenicol (for genomic-integrated biosensor strain, only 100 μg/mL ampicillin was added). A 2 μL aliquot of the overnight culture was inoculated into 198 μL OnEx system 2 medium supplemented with the appropriate antibiotics in a 96-well culture plate. The cultures were incubated at 30 °C with shaking at 700 rpm in a thermomixer for 48 hours. After 48 h cultivation, 10 μL of each culture was further diluted into 190 μL OnEx system 2 medium in 96 well round-bottom plate (Cat#262162, Nunc). All wells were analysed by GUAVA easyCyte Flow Cytometer and 5000 cells from each well were counted. All strains were inoculated in triplicate.

### Flow cytometry analysis and cell sorting by SONY SH800S Cell Sorter

For the flow cytometry analysis of *E.coli* BL21(DE3) G-TPAsensor (locus 4 & 5), cells were cultivated in OnEx system 2 medium supplemented with 0, 1 and 5 mM Na_2_TPA at 37 ℃ for 48 h. Each sample has triplicates. After cultivation, cells were diluted with sterilised PBS and 5000 events per sample were analysed by SONY SH800S Cell Sorter.

For mutant library analysis and sorting, after droplets cultivation and de-emulsification, the cells were diluted with PBS (pH 7.0) to achieve a flow cytometric event rate of approximately 3000 events/second in a SONY SH800S Cell sorter (SONY Biotechnology) equipped with a 70 μm nozzle size sorting chip. Prior to sorting, the cell suspension was filtered through a 35 μm cell strainer. The cell sorter was operated at pressure level 4, utilising 488 nm excitation lasers, with emissions measured through the FL1 filter for sfGFP fluorescence. A total of 10,000 cells from different mutagenesis libraries, along with a positive control group (*E. coli* BL21(DE3) G-TPAsensor (locus 4 & 5) with pET21b-*Is*PETase plasmid) and a negative control group (*E. coli* BL21(DE3) G-TPAsensor (locus 4 & 5) with pET21b blank plasmid), were initially analysed by plotting forward scatter area (FSC-A) against fluorescence. Groups whose fluorescence histograms showed no left shift compared to the positive control (0.1 and 0.2 mM MnCl2 groups) were pooled and subjected to the sorting procedure..

The flow cytometric event rate was maintained at ∼3000 events/second using the SONY SH800S cell sorter with a 70 μm nozzle sorting chip. Cells were firstly gated by FSC-A and side scatter area (SSC-A), and the gated cells were then further analysed for GFP fluorescence (488 nm excitation), and cells with the fluorescence intensity were sorted onto agar plates in a 96-well format. FACS data were processed using FlowJo software v10.8 (TreeStar). The sorted colonies were recovered in LB medium containing 100 μg/mL ampicillin, inoculated separately, and then transferred into induction medium for enzymatic assays or plasmid purification for sequencing.

### Protein expression and purification

PET degrading enzymes and variants were expressed in chemically competent *E.coli* BL21 (DE3). Single colonies from freshly transformed plates were cultured overnight at 37 °C in LB medium containing 100 µg/mL ampicillin. A 500 µL aliquot of the overnight culture was used to inoculate 50 mL of AIM TB medium (Cat#AIMTB0210, Formedium) supplemented with 100 µg/mL ampicillin. Cultures were grown at 37 °C with shaking at 200 rpm until reaching an OD600 of 0.8-1.0, then incubated at 16 °C for an additional 24 hours. Cell pellets were harvested by centrifugation at 5000 rpm for 20 minutes and then fully suspended in equilibration / binding buffer (pH 7.4, 50 mM NaH_2_PO_4_, 300 mM NaCl). Cell suspension was sonicated with 10% amplitude with 1s On / 1s Off for 10-20 min. The lysates were then centrifuged at 10,000 g for 20 minutes to separate the supernatant from the cell debris. The soluble fraction was subjected to affinity chromatography using Talon Metal Affinity Resin (Cat# 635504, Takara). Unbound proteins were washed off with 20 resin-bed volumes of equilibration buffer and 5 resin-bed volumes of wash buffer (pH 7.4, 50 mM NaH_2_PO_4_, 300 mM NaCl, 10 mM imidazole). Bound proteins were eluted using an elution buffer (pH 7.4, 50 mM NaH_2_PO_4_, 300 mM NaCl, 250 mM imidazole). The eluted proteins were desalted using a Hitrap Desalting Column (Cat#GE17-1408-01, Merck) and eluted in binding buffer (pH 7.4, 50 mM NaH_2_PO_4_, 300 mM NaCl). Protein purity was analysed by SDS-PAGE, and concentrations were measured using A280 with a Nanodrop (ThermoFisher).

For SDS and western blot, samples were prepared by mixing with 4x NuPage LDS sample buffer (Cat#NP0007, ThermoFisher) and boiling for 10-minute boiling at 85°C. Subsequently, 10 μL samples, along with a Spectra Multicolor Broad Range Protein Ladder (Cat#26634, ThermoFisher) were loaded onto 4–20% Mini-PROTEAN TGX stain-free protein gels (Cat#4568096, Biorad) for electrophoresis, which was run at 180 V for 40 minutes. Following electrophoresis, proteins were transferred to Trans-Blot Turbo Mini PVDF membranes (Cat#1704156, Biorad). The membrane was blocked for 2 hours at room temperature in PBST (PBS + 0.1% Tween 20) with 5% milk, washed with PBST for 3 times (5 min each), and incubated for 1 h at room temperature with 6x-His tag monoclonal antibody (HIS.H8) (Cat#MA1-21315, ThermoFisher). After three more PBST washes, the membrane was incubated for 1 h at room temperature with HRP conjugated anti-mouse IgG (H+L) secondary antibodies (Cat#31430, ThermoFisher), and washed again with PBST for 3 times (5 min each), and incubated for 5 min with West Pico plus HRP substrate (Cat#34580, ThermoFisher). Blotted proteins were visualised with a ChemiDoc XRS image analyzer (Bio-Rad).

### HPLC analysis

UPLC analysis was performed on a Thermo Fisher Dionex UHPLC-PDA-FLD LC system, equipped with an autosampler and a PDA detector set to 240 nm. The analysis utilised a Zorbax Eclipse Plus C18 Column (Agilent) with a stepped gradient solvent ratio method. Mobile phase A consisted of water with 20 mM phosphoric acid, while mobile phase B was methanol, with a constant flow rate of 1.2 mL/min. A 10 µL sample was injected for analysis.

Following injection, the mobile phase was initially set to 20% buffer B, then gradually increased to 65% over 7.0 minutes to separate TPA and MHET. The gradient was then rapidly decreased back to 20% within 6 seconds, followed by post-equilibration at 20% for 0.9 minutes, resulting in a total run time of 8.0 minutes. Peaks were identified by comparing them to chemical standards prepared from commercial TPA (Cat#40818, Sigma) and MHET (Cat#BD00777875, BLDPharma). Peak areas were integrated using Dionex Chromeleon software. With this method, TPA eluted at approximately 4.72 minutes, MHET at around 5.18 minutes, and small amounts of bis(2-hydroxyethyl) terephthalate (BHET) at approximately 5.80 minutes. Standards of TPA and MHET were analysed in each batch. TPA and MHET concentrations in the samples were estimated using standard curves (Supplementary Figs. 11a, 11b shows the TPA and MHET standard curves from one batch).

### Enzymatic assay

The sorted mutants underwent two additional rounds of assay screening. The first round was a supernatant assay. After overnight culturing in LB medium, cells were cultured in OnEx System 2 medium supplemented with 100 μg/mL ampicillin and 0.2 g/L PET nanoparticles at 30°C and 700 rpm for 48 hours, with all samples in triplicate. Following cultivation, the cells were centrifuged, and the supernatant was subjected to HPLC analysis The second round involved a purified enzyme assay. Variants that showed enhanced activity compared to the original *Is*PETase in the supernatant assay were further purified and quantified using the previously described protein purification methods. For the second round of screening, 40 nM of purified enzymes were thoroughly mixed with 4 g/L PET nanoparticles in 100 mM KH_2_PO_4_-NaOH buffer (pH 8) and incubated at 30°C with shaking at 700 rpm for 48 hours. The total assay volume was 600 μL in a 1.5 mL Eppendorf tube.

For additional enzymatic assays, 200 nM enzymes were typically used in a 600 μL buffered system in a 1.5 mL Eppendorf tube, with the tubes placed in a thermomixer set to the desired temperature and shaken vigorously at 700 rpm, unless otherwise specified. The buffers used in the assays included 100 mM KH_2_PO_4_-NaOH (pH 8.0) and 100 mM glycine-NaOH (pH 9.0). For assays conducted at different pH levels, buffers were prepared as follows: pH 6.0, 6.5, 7.0, 7.5, and 8.0 were prepared with 100 mM KH_2_PO_4_ and NaOH, while pH 9.0, 9.5, and 10.0 were prepared with 100 mM glycine and NaOH.

For the preparation of PET substrates, PET nanoparticles were washed 2-3 times with the specified buffers and then suspended to achieve the desired concentrations. PET products were punched into 6 mm-diameter discs, then washed sequentially with 1% SDS, 70% ethanol, and deionized water before use.

### Enzyme Kinetics

Around 1.5 mg PET disc (equivalent to a quarter of a 6 mm diameter-sized PET disc from the postconsumer yoghourt container lid) was added into 125 μl reaction solution in PCR tubes. The reaction solution consisted of 100 mM KH_2_PO_4_・NaOH pH8 buffer and varying concentrations of enzymes (5, 10, 25, 50, 75, 100, 150, 200, 250, 300, 400, 500 nM). The PCR tubes were incubated in a Biorad PCR machine set at 50 °C for both the block and the lid. Each enzyme concentration was tested in triplicate. A 3 μL sample of the reaction solution was taken at 1, 2, 3, 4, and 7 hours for measurement at 260 nm using a NanoDrop, followed by a UV light absorbance method. If necessary, samples were diluted with the reaction buffer. The values reported by the NanoDrop 1000 are based on a path length of 1 cm. Standard curves for TPA and MHET were also generated (Supplementary Figs. 11c, 11d), and a weighted mean coefficient of 965 M^-1^ cm^-1^, corresponding to a combination of 30% TPA and 70% MHET, was used. Reaction velocities at different enzyme concentrations were calculated by linear fitting. Inverse MM kinetics of Fast_2.9, FastPETase and LCC^ICCG^ data was fitted using the Inverse Michaelis–Menten model, which is 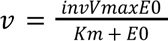^87^.

### Post Consumer PET degradation in 15 ml Bioreactor

Approximately 180 mg of post-consumer PET (yoghourt container lids, as mentioned above) was cut into small pieces (<1 cm x 1 cm) and added to a 20 mL bioreactor (Pioreactor 20 mL, Version 1.0, Pioreactor, 2024. Available at: https://pioreactor.com. Accessed on April 18, 2024) containing 15 mL of 100 mM KH_2_PO_4_-NaOH buffer (pH 8.0) and 100 nM of purified enzyme (0.24 mg enzyme/g PET) Fast_2.9, FastPETase, or LCC^ICCG^. Each sample was prepared in triplicate. The bioreactors were maintained at 50 °C with a stirring rate of 800 rpm, and the pH was regulated between 8.00 and 8.40 every 12 hours.

Samples of 100 µL were taken at various time points for time course analysis. Each sample was diluted to fall within the linear detection range of TPA and MHET for HPLC analysis.

Depolymerization was calculated based on the produced TPA equivalents (TPAeq) using the following formula:

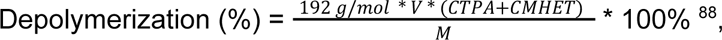

where 192 g/mol is the mass per repeating unit of PET, C_TPA_ and C_MHET_ are the molar concentrations of TPA and MHET as measured by HPLC, and M is the initial mass weight, approximately 180 mg, used in the Pioreactor.

### Coloured PET degradation

A piece of 40.2 mg green PET disc (from post consumer meat container product) was added into 3 mL 100 mM KH_2_PO_4_ ・NaOH (pH 8.0) buffer containing 0.0096 mg Fast_PETase (equal to 0.24 mg_enzyme_・g ^-1^) in a glass vial. The experiment was conducted in triplicate. The vials were placed in an incubator set at 50°C. An additional 0.0096 mg of Fast_2.9 enzyme was supplemented to the system every 24 hours, and the pH was adjusted to ∼8.0 as needed.

Samples of 10-50 µL were taken at various time points for time course analysis. Each sample was diluted to fall within the linear detection range of TPA and MHET for HPLC analysis. Images were captured at 2-minute intervals and compiled into a video.

### Polyester textile pretreatment and degradation by Fast_2.9

Approximately 200 g of each of the three types of polyester textiles (the outer skirt of a pink dress, the inner skirt of a pink dress, and a yellow T-shirt) were extruded using a capillary rheometry set at 280 °C, and immediately quenched in ice-cold water. The extruded material was then ground into powder by a cryomil.

Approximately 7-8 mg (∼12 g/L) pretreated polyester textile material was added into 100 mM KH_2_PO_4_ ・NaOH (pH 8.0) buffer with 100 nM (0.24 mg_enzyme_・g ^-1^) purified Fast_2.9 enzyme. An additional 100 nM of purified Fast_2.9 enzyme was supplemented every 24 hours, and the pH was adjusted to ∼8.0 using 5M NaOH every 24 hours. Samples of 50 μL were taken at 24, 48, and 72 hours for monomer content measurement.

Depolymerization was calculated based on the produced TPA equivalents (TPAeq) using the following formula:

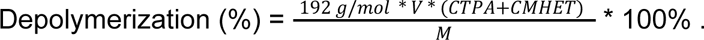

At the end of the 72 hours, the residual textiles were weighed using lyophilization to measure weight loss. The overall weight loss after degradation was also calculated by comparing it to the initial material weight.

### Structural analysis and molecular dynamics modelling

The starting coordinates for the *Is*PETase simulations were obtained from the crystal structure with PDB code 6EQE ^53^ and for FastPETase from the crystal structure with PDB code 7SH6 ^18^. The PDB files were pre-processed with the dock-prep tool of Chimera (v1.16)^89^, to complete incomplete side chains (using the Dunbrack 2010 library) and remove the low occupancy alternative locations of side chains when present. Residue S269 was mutated with the mutate_model.py script of MODELLER (v10.4)^90^ to T269. For the wild type, S269 was maintained as S269.

System parameterization was performed using the CHARMM-GUI web interface ^91,92^. Hydrogens were assigned based on physiological pH, and the systems were placed on a rectangular water box of TIP3P ^93^ water molecules extending at least 10 Å from any protein atom. The solvated systems were neutralised using Cl- ions and an additional 150mM NaCl were added. The CHARMM36m force field was used to parametrize the protein ^94^.

The systems were first minimised for a maximum of 5000 steps, using the steepest descent algorithm for the first 2500 steps and then switching to the conjugate gradient method. Weak positional restraints of 1 kcal mol^-1^ angstrom^-2^ were added to all the protein atoms. Following minimization, the systems were equilibrated under NVT ensemble for 125 ps, during which the temperature was raised to 303.15K for the IsPETase simulations and 313K for the FastPETase ones. Positional restraints were maintained during equilibration, with an integration time step of 1 fs.

All production simulations were conducted under the isothermal, isobaric conditions (NPT) with 1 bar pressure, using the Langevin Thermostat ^95^ (friction coefficient 1 ps-1) and Montecarlo Barostat ^96^. Periodic boundary conditions and Particle Mesh Ewald (real space cutoff 12 Å and grid spacing 1.2 Å) were employed for remote electrostatic interactions ^97^, while Van der Waals contacts were truncated at the real space cutoff. Protein bonds involving hydrogens were constrained using the SHAKE algorithm, and water molecules were treated with SETTLE, allowing for an integration time step of 2 fs.

Simulations were run using the single precision pmemd.cuda implementation of Amber 2022^98^. Each system was simulated for a total of 1 µs, with simulation coordinates saved periodically for further analysis. The initial 500 ns were discarded as the equilibration phase.

Root mean square fluctuation (RMSF) profiles for each system were obtained using cpptraj ^98^. Trajectory frames were aligned using the backbone atoms (C, CA, N), and residue weighted RMFS profiles were generated using atomic masses as weights. To compare systems, the area under the curve (AUC) of the RMSF profile from residues 88 to 91 was calculated, and significance was assessed using a two-tailed independent t-test.

Root mean square deviations (RMSD) profiles for residues 88 to 91, which showed significant fluctuations between the S269 and S269T variants, was computed using the mdtraj python package ^99^. Frames were aligned to the crystal structure using the backbone atoms and excluding the region of residues of interest. The RMSD was computed for C-α atoms from residues 88 to 91 without realigning the trajectory to better capture different conformations. Results from the different replicates were combined, and the probability density was computed with scipy using the default number estimation of bins. Representative structures for each of the RMSD peaks of the residues 88 to 91 were determined by randomly sampling 50 snapshots, computing an average structure, aligning it to the docking structure, and visualising it in PyMOL (http://www.pymol.org/pymol).

Correlation paths were computed using WISP ^55^ by combining all the replicates of *Is*PETase S269T. Paths were calculated using residue T269 as the source residue and residues T88 to S93 as the sink, extending until reaching residue T88. Simulations were converted to PDB format and saved 10 times less frequently. The top 10 shortest paths were computed using default WISP parameters and visualised in PyMOL, with path thickness representing the frequency of occurrence..

## Data and Code Availability

Data supporting the findings of this study are available within the paper and its Supplementary Information.

## Supplementary information

Supplementary Methods, Supplementary Figs. 1-11, Supplementary Tables 1-6, Supplementary Note, Supplementary video

## Acknowledgements

The study was supported by SciLifeLab fellows program (A.Z.), Swedish Research council (Vetenskapsrådet) starting grant no. 2019-05356 (A.Z.), Formas early-career research grant 2019-01403 (A.Z.), WALP Wallenberg Launchpad project 2023.0523 supported by the Knut and Alice Wallenberg Foundation (A.Z.). A.Z. is a Marius Jakulis Jason Foundation scholar. We thank Roland Kádár and Sajjad Pashazadehgaznagh from Chalmers University of Technology for their help with textile pretreatment. We acknowledge Jacob Kindbom from Chalmers Mass Spectrometry Infrastructure for providing training and instruction on HPLC usage, Stefan Gustafsson from Chalmers Materials Analysis Laboratory on SEM training and Katarina Logg on FT-IR guidance. We thank Shuichi Haraguchi from Chalmers University of Technology for discussions on DSC results. We thank Mukil Madhusudanan from Technical University of Denmark for discussions on nanoparticles. We thank Shanjun Ruan for videography and video editing.

