## Supplementary Information for "Single Distal Mutation Enhances Activity of known PETases via Stabilisation of PET-binding"

### Supplementary Methods

#### ρ-nitrophenyl acetate (ρNP-Ac) assay

A 20 mM solution of ρ-nitrophenol acetate (ρNP-Ac) (Cat#N8130, Sigma) was prepared by dissolving 7.246 mg of ρNP-Ac in 2 mL of 96% ethanol and kept on ice until use. The assay was performed in a Nunc 96-well transparent plate by mixing 100 µL $NaH_2PO_4$ buffer (pH 7.0), 80 µL Milli-Q water, 10 µL cell culture supernatant, and 20 µL of the 20 mM ρNP-Ac solution. The cell cultures were grown overnight in Overnight Express OnEx system 2 (Cat#71300, Merck), and the supernatant was collected after centrifugation at 10,000 rpm for 10 minutes.

Absorbance at wavelength 405 nm was measured every 30 s for 20 min with plate reader (SPECTROstar Nano, BMG Labtech), which was pre-set at 30 °C. Conversion from absorption values to the catalytic product ρ-nitrophenoxide concentration was carried out using Beer-Lambert law c=A/(d*ε), in which an extinction coefficient ε value is 12.8 $mM^{-1} \cdot cm^{-1}$, and path length for 200 µl in 96-well plate (d) is 0.335 cm.

#### Western blotting

For cell lysate preparation, 500 µL of cells were collected by centrifugation, washed with 500 µL Milli-Q water, resuspended in 100 µL Milli-Q water, and heated at 100 °C for 15 minutes. A 5 µL sample was mixed with 1 µL Milli-Q water, 1 µL 1 M dithiothreitol (DTT), and 2.5 µL 4x loading dye, then heated at 100 °C for 15 minutes. For Click-PEGylation samples, DTT was replaced with Milli-Q water. Equal amounts of protein were separated by 4-15% SDS-PAGE gel (Cat#4568084, Bio-Rad) and transferred onto PVDF membranes (Cat#1704274, Bio-Rad).
Western blotting was performed using a 6x-His tag monoclonal antibody (Cat#MA1-21315, ThermoFisher) and the Pierce Fast Western Blot Kit (Cat#35055, ThermoFisher) according to the manufacturer's instructions.

#### Size measurement of PET nanoparticles

PET NP were suspended in milliQ water and sonicated for 10 min before Multi-angle dynamic light scattering (MADLS) analysis. The hydrodynamic diameter of the NPs was measured using the Malvern zetasizer ultra (Malvern Panalytical).

#### Fourier-transform infrared spectroscopy (FT-IR) analysis

Prior to the analysis, the PET material was subjected to washing with 1% sodium dodecyl sulfate (SDS), ethanol, and water once each, followed by drying. PET NP was washed with Milli-Q water three times and subsequently lyophilized to obtain PET NP powder.
ATR-FTIR measurements of PET were performed with a Vertex 70v spectrometer (Bruker Optics) coupled with a Hyperion 3000 microscope (Bruker Optics) equipped with a germanium (Ge)-attenuated total reflectance objective lens (ATR 20×) and a liquid nitrogen cooled mercury cadmium telluride (MCT) detector. The spectra were obtained from 800 to 4000 $cm^{-1}$ at a resolution of 4 $cm^{-1}$ with 32 repeated scans. Background scanning was performed before sample loading each time.

### PET crystallinity analysis by DSC

DSC was used to determine the percentage crystallinity of the PET materials. PET film samples (3-5 mg) were placed in aluminum Tzero pans with a Tzero solid sample lid. Samples were first heated from 40 to 300 °C at 5 °C min, held at 300 °C for 1 min, cooled from 300 to 30 °C at –5 °C min−1 and held at 30 °C for 1 min in a DSC250 (TA Instruments) with a RCS90 electric chiller. The percentage crystallinity was determined on the first heating scan using the enthalpies of melting and cold crystallization. The equation used to calculate percentage crystallinity within the PET film was the following

$$Xc = \frac{\Delta Hm - \Delta Hcc}{\Delta Hf100} \times 100\%$$

Where *Xc* = Degree of crystallinity

$\Delta H_m$ = Enthalpy of melting enthalpy

$\Delta H_{cc}$ = Enthalpy of cold crystallization enthalpy

$\Delta H_{f100}$ = Heat of fusion of 100% PET Crystallization = 140 J/g [1].

### Scanning electron microscopy (SEM)

SEM was performed using a Zeiss Ultra 55 FEG scanning electron microscope (FESEM) (Carl Zeiss Microscopy) at 5 kV accelerating voltage and a beam current of 10 µA. Prior to SEM analysis, all samples were sputter coated with 6 nm of gold palladium using a low vacuum sputter coater (Leica EM ACE200, Leica Microsystems Inc.).

### Click-PEGylation assay

All purified proteins (*Is*PETase, M2.9, ThermoPETase. ThermoPETase_2.9, FastPETase, FastPETase_2.9) were normalized to the same concentration of 160 ng/µl by buffer exchange with 20 mM sodium phosphate, 50 mM NaCl, pH 6.5 buffer in Amicon Ultra 0.5 mL filters (10 kDa cutoff, Cat#UFC5010, Merck). Approximately 2.8 µg of each proteins were incubated with 5 mM methoxypolyethylene glycol maleimide (NEM-mPEG, Cat#63187, Sigma) and 3% SDS, and then heated for 10 min at 65 °C. Negative controls were incubated with 10 mM N-ethylmaleimide (NEM, Cat#E1271, Sigma) and 3% SDS under the same conditions. After incubation, results were analyzed by western blotting as described above.

### Supplementary Figures

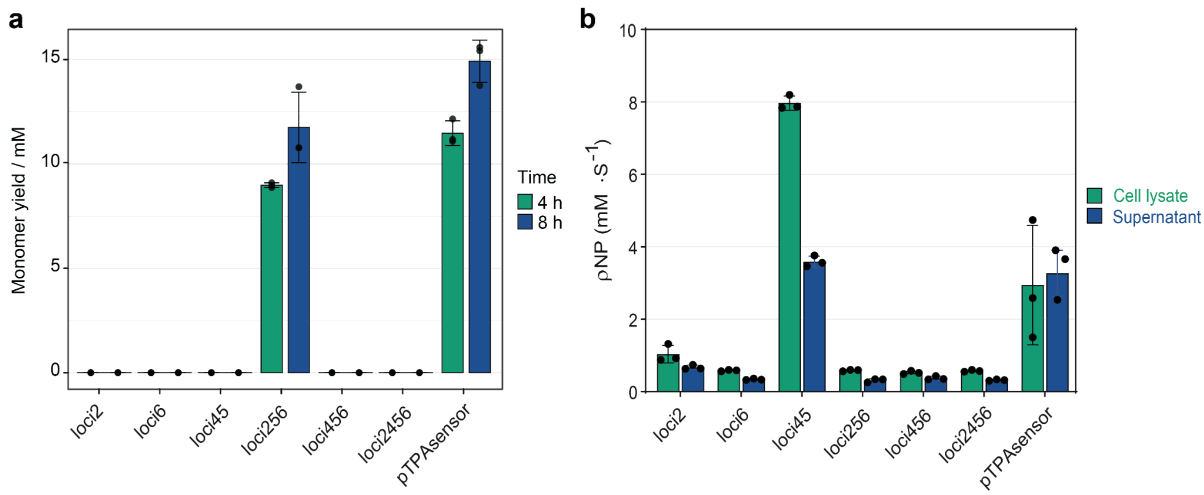

**Supplementary Figure 1. a**, Total monomer (TPA and MHET) yield after incubating both the supernatant and cell lysate of engineered *E. coli* BL21(DE3) strains with genomic TPAsensor insertion and *E. coli* BL21(DE3) with the pTPAsensor plasmid (pTPAsensor) with PET nanoparticles (PET NP) for 4 and 8 hours at 30 °C. **b**, ρNP consumption rate after incubating the supernatant and cell lysate of the same strains with 20 mM ρNP-Ac for 30 minutes at 30 °C. All data are shown as mean ± SD (n=3).

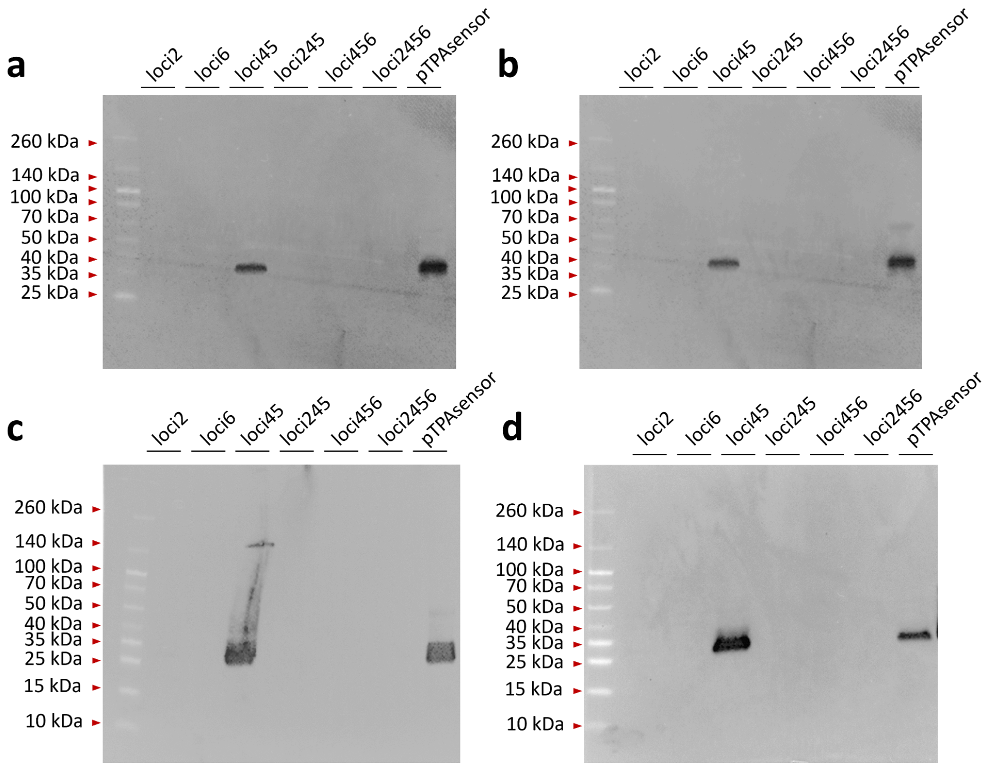

**Supplementary Figure 2.** Western blotting analysis of *Is*PETase overexpressed from engineered *E. coli* BL21(DE3) strains with genomic TPAsensor insertion and *E. coli* BL21(DE3) with the pTPAsensor plasmid (pTPAsensor). Both cell lysates **(a, c)** and supernatant **(b, d)** samples were analyzed for strains cultivated in IPTG-supplemented induction medium **(a, b)** or autoinduction medium **(c, d)**.

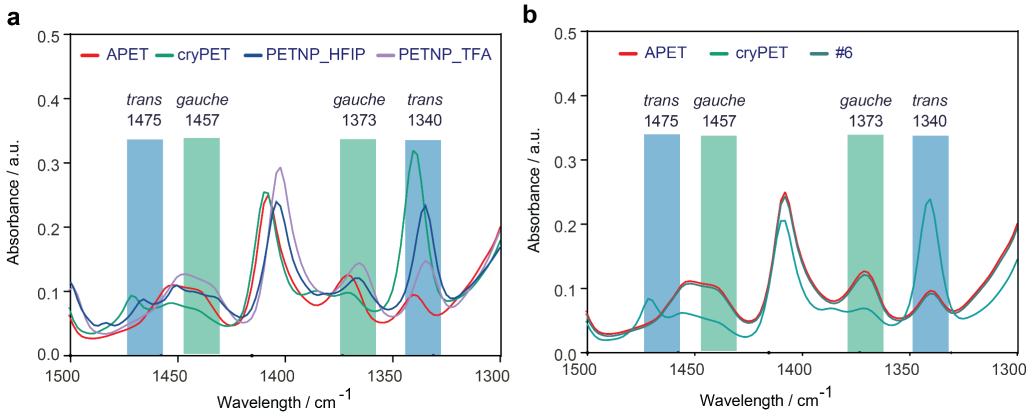

**Supplementary Figure 3.** FT-IR analysis of PET material. **a,** Amorphous PET film from GoodFellow (APET), high crystalline PET film from GoodFellow (cryPET), PET NP prepared from amorphous PET film with solvent HFIP (PETNP_HFIP) and PET nanoparticles prepared from amorphous PET film with solvent TFA (PETNP_TFA). **b,** Amorphous PET film from GoodFellow (APET), high crystalline PET film from GoodFellow (cryPET) and the PET film from the postconsumer yogurt container lid (#6).

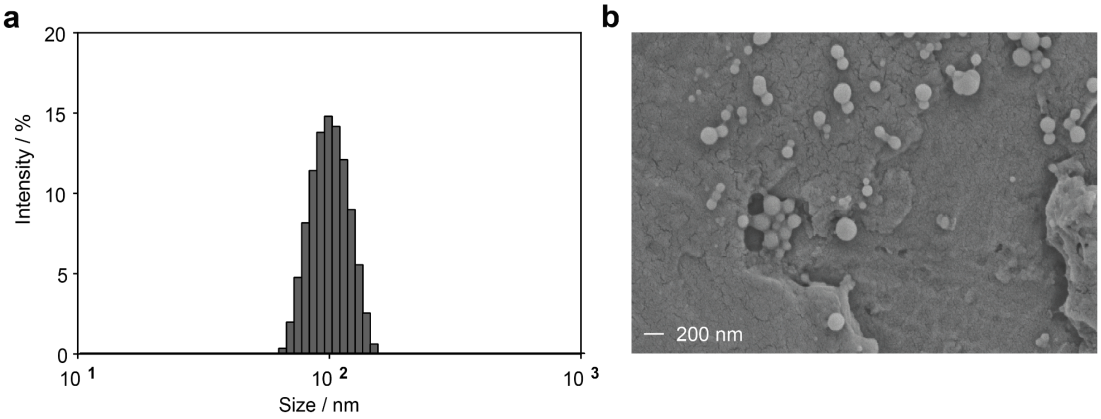

**Supplementary Figure 4. Size characterization of homemade PET nanoparticles with HFIP (PETNP_HFIP). a,** The size distribution of PET nanoparticles analyzed by MADLS. **b,** The PET nanoparticles analyzed by SEM. The SEM pictures were taken twice, with the pictures shown being representative of the set.

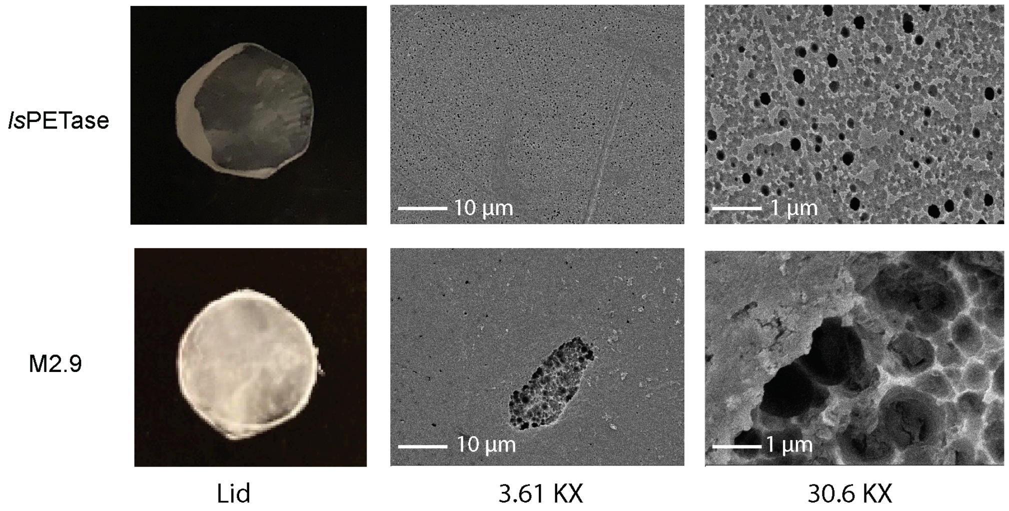

**Supplementary Figure 5.** Pictures and SEM analysis of the PET discs from the postconsumer yogurt contain lid, by 200 nM *Is*PETase and M2.9 in 100 mM $KH_2PO_4$ • NaOH (pH 8.0) buffer at 50 °C for 48 h. Photos were taken twice, with the photos shown being representative of the set.

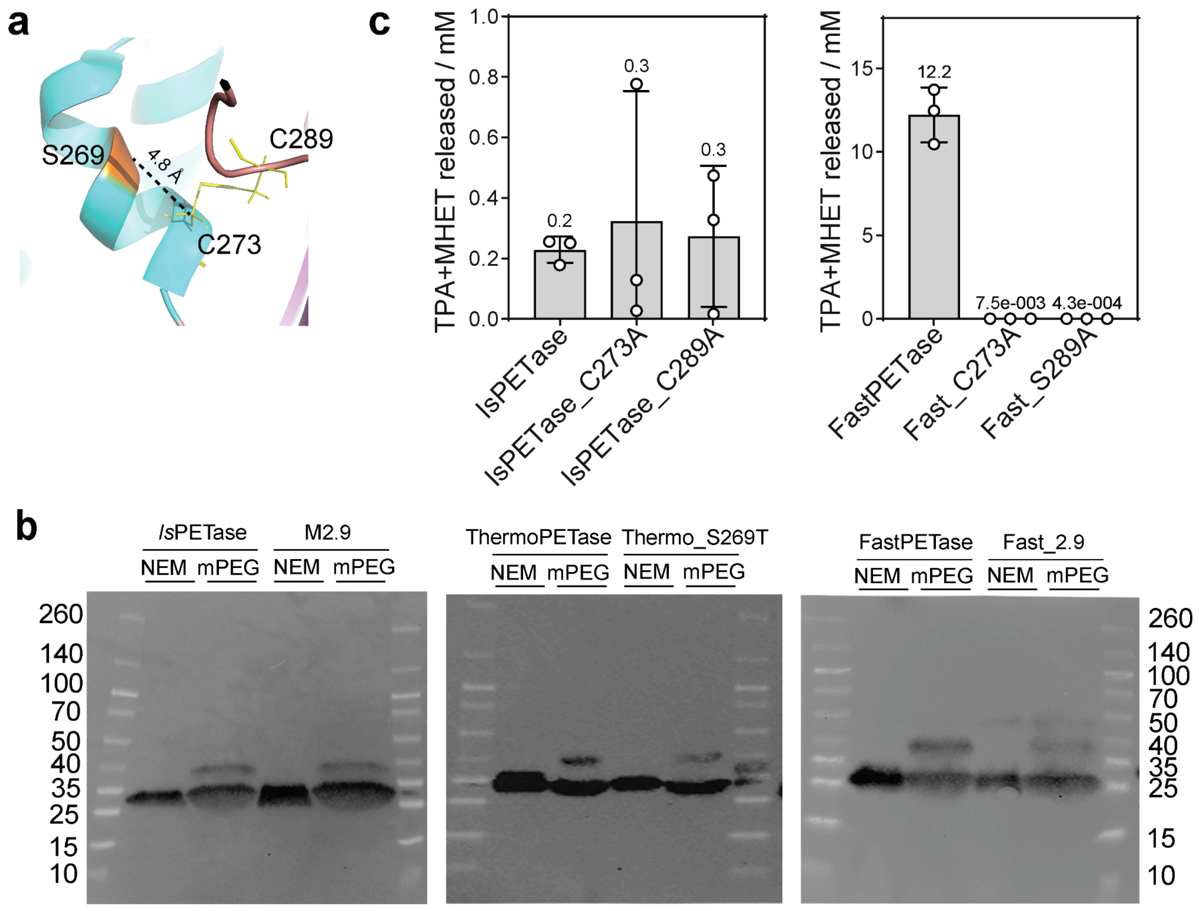

**Supplementary Figure 6. a,** Structural position of S269, C273, C289 in IsPETase. Dash line indicates the distance between S269 and C273 is 4.8 Å. **b**, Click-PEGylation analysis of functional *Is*PETase and M2.9, ThermoPETase and Thermo_S269T, FastPETase and

Fast_2.9. **c,** Released monomers by *Is*PEase, *Is*PEase_C273A, *Is*PETase_C289A and FastPEase, FastPEase_C273A, FastPETase_C289A. 200 nM enzymes were incubated with 6 mm diameter-sized postconsumer PET disc for 48 h and the released monomers were measured by HPLC. Each sample has triplicates and all data are shown as mean ± SD. The mean values of released monomers for each sample were added on top of each column.

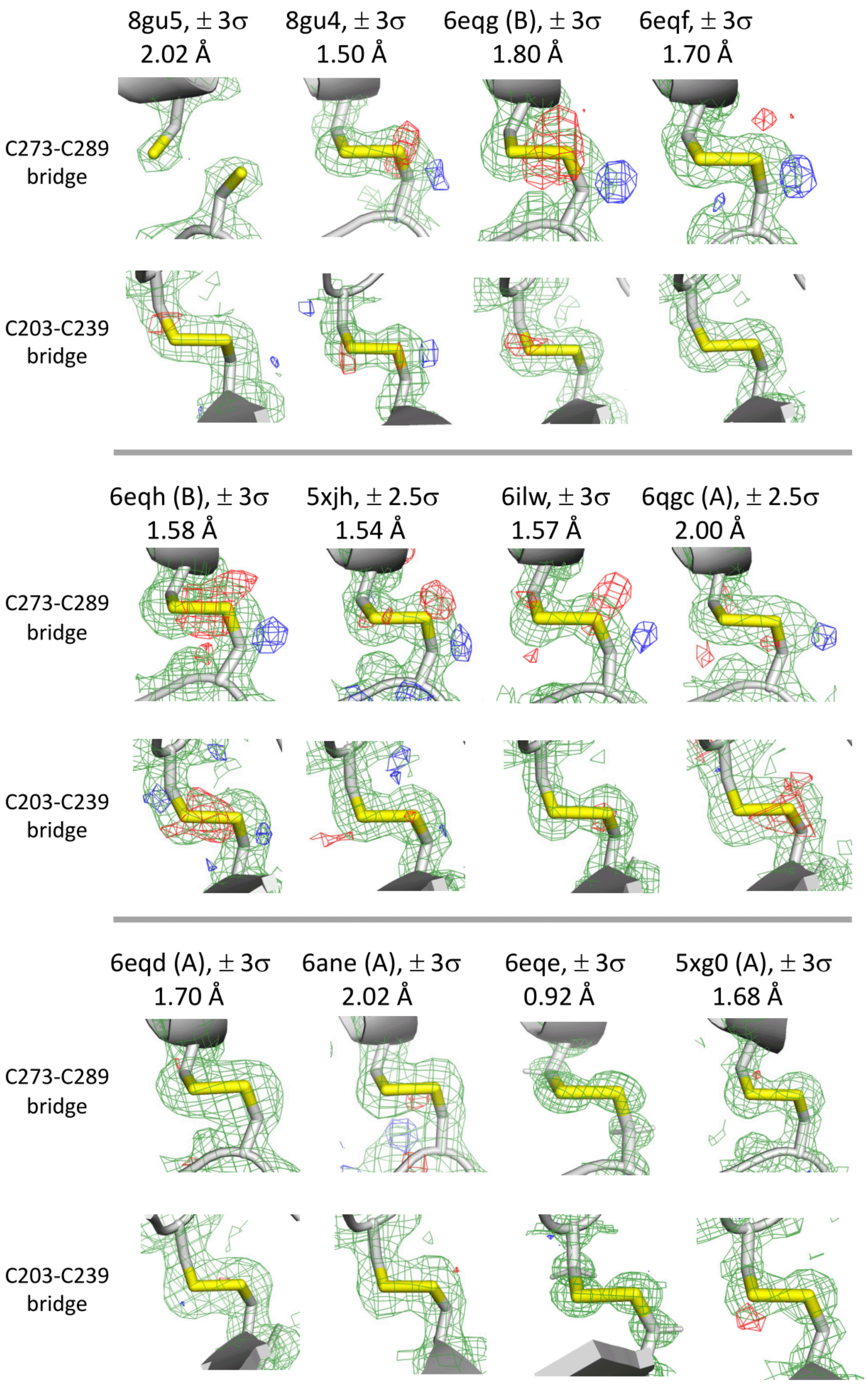

**Supplementary Figure 7.** Electron density maps of the disulfide-bridges of all wild type IsPETase entries in the PDB. The resolution of each structure and the chain that was used is indicated above the maps. For each structure, the 2Fo-Fc map at 1σ is indicated with green mesh; and the Fo-Fc maps at ± 3 or ± 2.5σ are indicated with blue (+) and red (-) mesh on each panel. For the bridge between C273-C289, in 7 out of 12 structures there is an indication of an alternative position of the sulphur atom of C289 (blue mesh), while in PDB ID: 8GU5 the CYS273 and CYS289 do not form a bridge at all. In contrast, only 2 structures indicate an alternative location of a sulphur atom in the C203-C239 bridge, and none of the bridges are open.

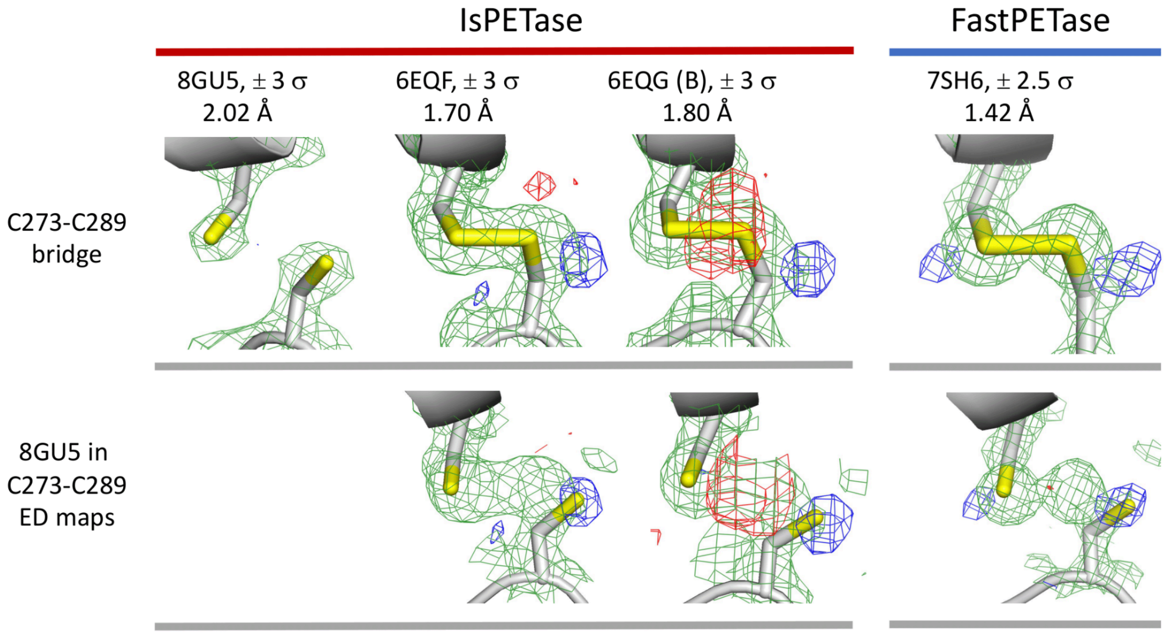

**Supplementary Figure 8.** Overlap between the CYS273 and CYS289 atoms of PDB ID 8GU5, and the electron density maps of selected structures (upper row). The location of the sulphur atom of CYS289 in 8GU5 largely overlaps with the additional peak (+3 σ, blue) that is frequently observed in many structures, both in IsPETase and FastPETase. Upper row: electron density maps of selected structures, Lower row: atoms of 8GU5 placed in the electron density maps of other structures.

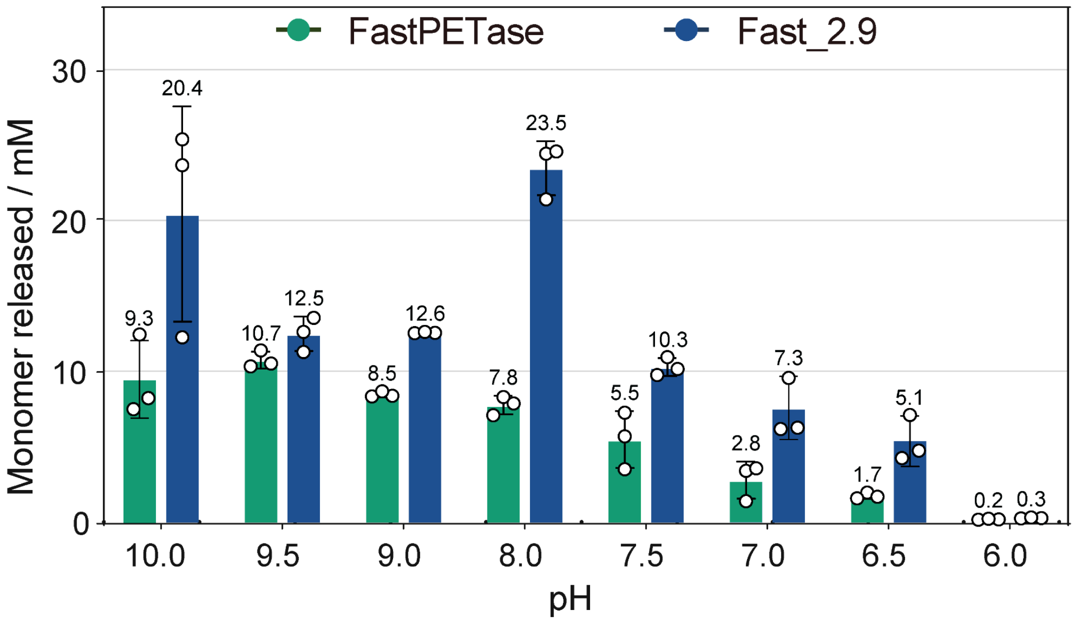

**Supplementary Figure 9.** Monomer release after incubating a 6 mm-diameter post-consumer PET disc with 200 nM FastPETase or Fast_2.9 in buffers of different pH at 50

°C for 48 hours (n=3). Data are presented as mean ± SD, with the mean values of released monomers indicated above each column.

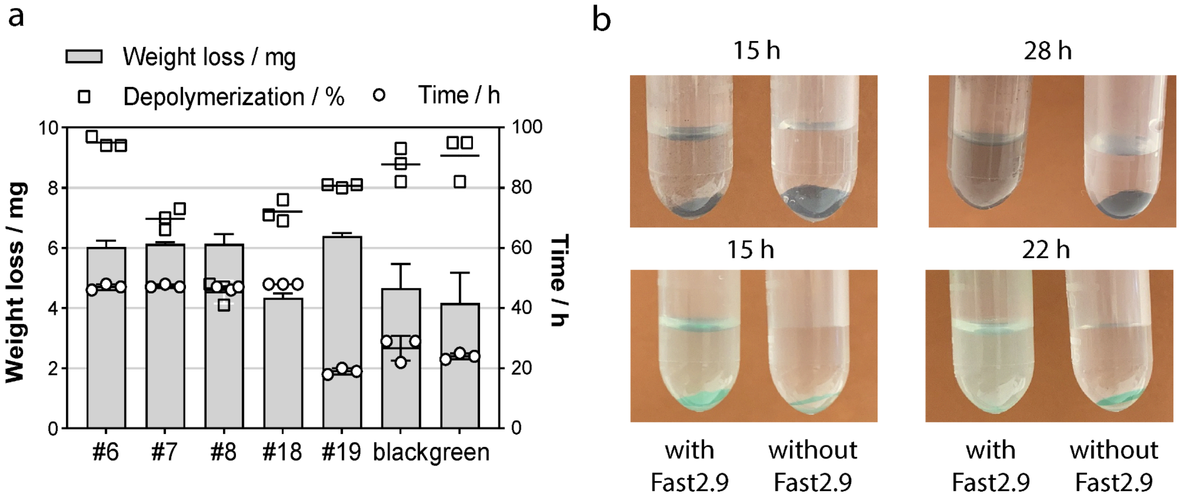

**Supplementary Figure 10. a,** Complete degradation of seven types of post-consumer PET products by Fast2.9 at 50°C, including two colored opaque PET containers. All samples were run in triplicate, and data are presented as mean ± SD. **b**, Images of green and black colored opaque PET at 15 hours and at the final time point of complete depolymerization. Photos were taken twice, with the displayed images being representative of the set.

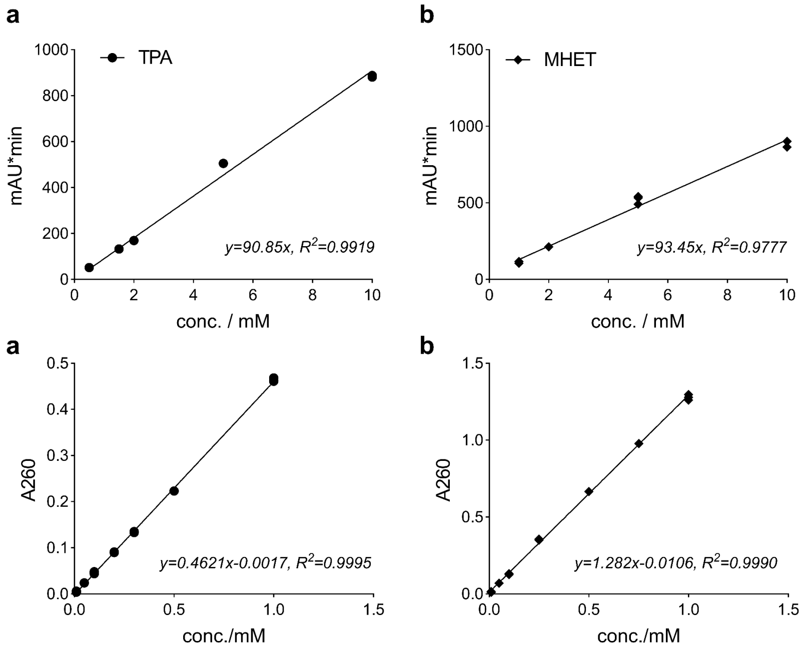

**Supplementary Figure 11.** An example of standard curves of TPA **(a, c)** and MHET **(b, d)** as measured by HPLC **(a, b)**, and by UV light absorbance method with Nanodrop **(c, d)** from one batch of experimetn. Each sample has triplicates at different concentrations.

### Supplementary Tables

**Table 1. TPA biosensor sequence**

| Sequence | Note |
|---|---|
| TTAGAGACCTTGCGGGTACAGCTTCTTCTCGAGTTCGTTGCG<br>CGCACGTTTCAGCGGGATCAGGAAGGTTTCTTTGAACTCGGA<br>CATGCTTAACCTTTCAGCGCGCACCGCGATCGACATTGCGGC<br>AATGGTATTACCTTGCGGATCACGCACCGGAGCTGCCATAGA<br>ACGAACGCCCAGCTCCAGTTCGCCGTCGGAACAAGACCAAC<br>CAGACTGACGGCATGTCTCGAGCAAGCCTAACAGTTCCTCCA<br>GATCGGTCACCGTATGAGGCGTCAGAGCAACACGTTCAATCA<br>TCTCCAAACGCGCACGCGCTTCTTGTTGCGGAAGGCCGGAC<br>AGCAGCATACGACCAATCGCGCTGCAGTAGACCGGTAAACG<br>GCTGCCAATACCCAGACCGGTACTAAGGCTACGGCGTGCGG<br>TGCTACGGCCGATGATAATGGCGTCGTCCTCAAGCAGGGTTC<br>CCAGGCTCGCGCTTTCACGGGTGCGTTCAGACAGCGCGTCC<br>AACAGCGGCTGTGCCAGCGCCGGCATCGGACGGCTCGACA<br>GGAAGCTATAGGCGATCAGCAGGCTTTTTGGCTGCATCCAAA<br>ACAGCTTACCATCGCTCTCCAGATAACCCAGCTGAACCAAGG<br>TGGACAAAGAACGTCTGGCGCTAGCCGGAGTGCTCTGGGTC<br>AAGCGCGCAACCTCGGACAGGGTCAGACGGGTGTGGCGAC<br>GATCAAAGCACGTCAGCACGCCCAGACCTTTGCGTAGTGATT<br>CTACAAAATTTTTATCTTGCAT*AGCGATCAAATCAAGGTGTTTT*<br>*CAACATTTTTGCGCATAGCGCAAAAACAGTGTACACCAAAGTA*<br>*CGACATCCTTACAATGCAGTTCCCCACACAAGAAGGAGATATA*<br>*CAT*ATGGTTAGCAAAGGTGAAGAACTGTTTACCGGCGTTGTG<br>CCGATTCTGGTGGAACTGGATGGTGATGTGAATGGCCATAAA<br>TTTAGCGTTCGTGGCGAAGGCGAAGGTGATGCGACCAACGG<br>TAAACTGACCCTGAAATTTATTTGCACCACCGGTAAACTGCCG<br>GTTCCGTGGCCGACCCTGGTGACCACCCTGACCTATGGCGT<br>TCAGTGCTTTAGCCGCTATCCGGATCATATGAAACGCCATGAT<br>TTCTTTAAAAGCGCGATGCCGGAAGGCTATGTGCAGGAACGT<br>ACCATTAGCTTCAAAGATGATGGCACCTATAAAACCCGTGCGG<br>AAGTTAAATTTGAAGGCGATACCCTGGTGAACCGCATTGAACT<br>GAAAGGTATTGATTTTAAAGAAGATGGCAACATTCTGGGTCAT<br>AAACTGGAATATAATTTCAACAGCCATAATGTGTATATTACCGC<br>CGATAAACAGAAAAATGGCATCAAAGCGAACTTTAAAATCCGT<br>CACAACGTGGAAGATGGTAGCGTGCAGCTGGCGGATCATTAT<br>CAGCAGAATACCCCGATTGGTGATGGCCCGGTGCTGCTGCC<br>GGATAATCATTATCTGAGCACCCAGAGCGTTCTGAGCAAAGAT<br>CCGAATGAAAAACGTGATCATATGGTGCTGCTGGAATTTGTTA<br>CCGCCGCGGGCATTACCCACGGTATGGATGAACTGTATAAAG<br>GCAGCtga | Underlined sequence at 5’ end is the transcription factor *tphR*, italic sequence is the intergenic region and underlined sequence at 3’ end is the *sfGFP*. |

**Table 2. The integrated loci of TPA biosensor in the BL21 (DE3) genome.**

| | Target locus | Gene | Protein | gRNA position in BL21 (DE3) genome (CP001509.3)_ | Spacer sequence (5'→3') |
|---|---|---|---|---|---|
| Loci 2 | aroG-gpmA locus | In between | | 745115-745146 | GTCAGAGTGGCGTATCCGATGAATCACCACAG |
| Loci 4 | purT-eda locus | *eda* | KHG/KDPG aldolase; 2-dehydro-3-deoxy-phosphogluconate/4-hydroxy-2- oxoglutarate aldolase | 1877854-1877885 | GCATTACTAAGCTGGCGCGTGAAGCTGTAGAA |
| Loci 5 | tktB-ypfG locus | *tktB* | transketolase 2, thiamine triphosphate-binding | 2446078-2446109 | GAGAATATTGTGGCAAAAGCGCATAAGGTGCT |
| Loci 6 | yghA-exbD locus | *yghA* | putative oxidoreductase | 3017942-3017973 | GCAGAAGTGCACGGCGTGTGCGGCGGCGAGCA |

**Table 3. Protein sequences**

| **Name** | **Protein sequence (His tag not included)** | **Note** |
|---|---|---|
| *Is*PETase_2.9 (*Is*PETase$^{S269T}$) | MNFPRASRLMQAAVLGGLMAVSAAATAQTNPYARGPNPTAASLEASAGPFTVRSFTVSRPSGYGAGTVYYPTNAGGTVGAIAIVPGYTARQSSIKWWGPRLASHGFVVITIDTNSTLDQPSSRSSQQMAALRQVASLNGTSSSPIYGKVDTARMGVMGWSMGGGGSLISAANNPSLKAAAPQAPWDSSTNFSSVTVPTLIFACENDSIAPVNSSALPIYDSMSRNAKQFLEINGGSHSCANSGNSNQALIGKKGVAWMKRFMDNDTRY**T**TFACENPNSTRVSDFRTANCS | The S269T mutation is highlighted in bold |
| FastPETase_2.9 (FastPETase$^{S269T}$) | MNFPRASRLMQAAVLGGLMAVSAAATAQTNPYARGPNPTAASLEASAGPFTVRSFTVSRPSGYGAGTVYYPTNAGGTVGAIAIVPGYTARQSSIKWWGPRLASHGFVVITIDTNSTLDQPESRSSQQMAALRQVASLNGTSSSPIYGKVDTARMGVMGWSMGGGGSLISAANNPSLKAAAPQAPWHSSTNFSSVTVPTLIFACENDSIAPVNSSALPIYDSMSQNAKQFLEIKGGSHSCANSGNSNQALIGKKGVAWMKRFMDNDTRY**T**TFACENPNSTAVSDFRTANCS | The S269T mutation is highlighted in bold |
| ThermoPETase_2.9 (ThermoPETase$^{S269T}$) | MRGPNPTAASLEASAGPFTVRSFTVSRPSGYGAGTVYYPTNAGGTVGAIAIVPGYTARQSSIKWWGPRLASHGFVVITIDTNSTLDQPESRSSQQMAALRQVASLNGTSSSPIYGKVDTARMGVMGWSMGGGGSLISAANNPSLKAAAPQAPWHSSTNFSSVTVPTLIFACENDSIAPVNSSALPIYDSMSRNAKQFLEINGGSHSCANSGNSNQALIGKKGVAWMKRFMDNDTRY**T**TFACENPNSTAVSDFRTANCSLEDPAANKARKEARLAAATAEQ | The S269T mutation is highlighted in bold |
| HotPETase_2.9 (HotPETase$^{S269T}$) | MQTNPYARGPNPTAASLEASAGPFTVRSFTVARPVGYGAGTVYYPTNAGGTVGAIAIVPGYTATQSSINWWGPRLASHGFVVITIDTNSTLDKPESRSSQQMAALRQVASLNGTSSSPIYGKVDTARGGVMGWSMGGGGSLISAANNPSLKAAVMAPWHSSTNFSSVTVPTLIFACENDRIAPVKEYALPIYDSMSLNAKQFLEICGGSHSCACSGNSNQALIGMKGVAWMKRFMDNDTRY**T**QFACENPNSTAVCDFRTANCS | The S269T mutation is highlighted in bold |
| DepoPETase_2.9 (DepoPETase$^{S269T}$) | MNFPRASRLMQAAVLGGLMAVSAAATAQTNPYARGPNPTAASLEASAGPFTVRSFTVSRPSGYGAGTVYYPTNAGGTVGAIAIVPGYIARQSSIKWWGPRLASHGFVVITIDTNSTLDQPSSRSSQQMAALRQVASLNGTSSSPIYGKVDTARMGVMGWSMGGGGSLISAANNPSLKAAAPQAPWHSSTNFSSVTVPTLIFACENDSIAPVNSSALPIYNSMSRNAKQFLEIKGGSHSCANSGNSDQALIGKKGVAWMKYFMDNDTRYS**T**FACENPNSTRVSDFRTANCPAAA | The S269T mutation is highlighted in bold |
| LCC$^{ICCG}$ | MSNPYQRGPNPTRSALTADGPFSVATYTVSRLSVSGFGGGVIYYPTGTSLTFGGIAMSPGYTADASSLAW | |

| | |
|---|---|
| | LGRRLASHGFVVLVINTNSRFDGPDSRASQLSAAL NYLRTSSPSAVRARLDANRLAVAGHSMGGGGTLRI AEQNPSLKAAVPLTPWHTDKTFNTSVPVLIVGAEA DTVAPVSQHAIPFYQNLPSTTPKVYVELCNASHIAP NSNNAAISVYTISWMKLWVDNDTRYRQFLCNVND PALCDFRTNNRHCQ |

**Table 4. Characterization and crystallinity of postconsumer PET materials used in this study**

| PET product | Picture | Crystallinity[1] | A1340/A1372[2] | A1475/A1457[3] |
|---|---|---|---|---|
| #6 yogurt container lid | 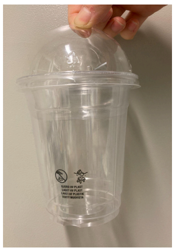 | 1.2% | 0.7924 | 0.4054 |
| #7 blueberry container | 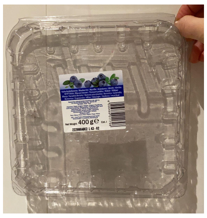 | 2.1% | 0.8128 | 0.4176 |
| #8 tomato container | 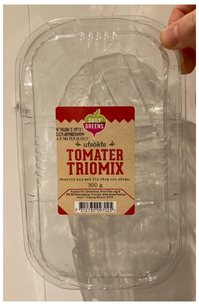  | 5.3% | 0.8399 | 0.4429 |
| #18 tips container | 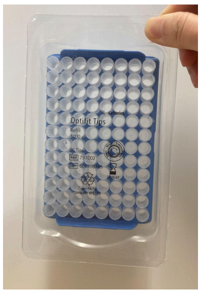 | 4.3% | 1.0982 | 0.4672 |
| #19 salad container lid | 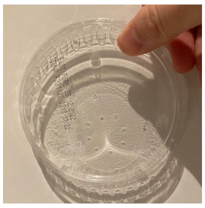 | 1.5% | 0.7947 | 0.4297 |

| | | | | |
|---|---|---|---|---|
| #black meat container | 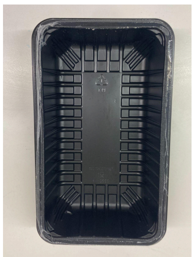 | 10.1% | 0.7390 | 0.4871 |
| #green meat container | 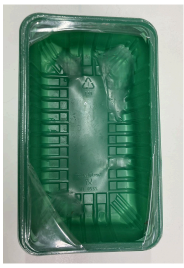 | 8.7% | 0.8715 | 0.6302 |
| GoodFellow_ amorphous PET (Cat#ES303010/19) | 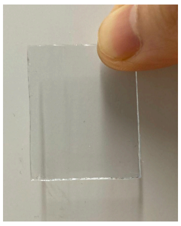 | 3.8% | 0.7839 | 0.4108 |
| GoodFellow_ crystalline PET (Cat#ES301230) | 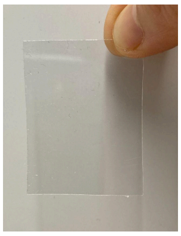 | 27.8% | 4.4245 | 0.8022 |
| PET NP_HFIP | | 4.2% | 0.9573 | 0.7933 |
| PET NP_TFA | | 10.3% | 1.4596 | 0.6353 |

[1] crystallinity was measured by DSC;
[2] The ratio of A1340 with A1372 as measured by FT-IR represents the ratio of *trans* to *gauche* conformations of glycol-CH2 wagging [6] ;

[3] The ratio of A1475 with A1457 as measured by FT-IR represents the ratio of *trans* to *gauche* conformations of glycol-$CH_2$ bending [6].

**Table 5. Purified variants and their mutations.**

| **Purified variants** | **Mutations** |
|---|---|
| M1.1 | L249P |
| M1.4 | Y63S |
| M1.5 | K227E/N233S |
| M1.6 | D186A |
| M1.8 | Q28H/K227Q |
| M1.9 | G66E |
| M1.10 | K227E |
| M2.1 | S278R |
| M2.3 | L15R/N30D/V107A/S207N |
| M2.4 | F57Y/D118G |
| M2.8 | S213G/D263G/T286A |
| M2.9 | S269T |

**Table 6. Primers used in this study**

| Primer name | Primer sequence |
|---|---|
| PETase_RM_SP_F | CAGCGGCCACCGCGCAG |
| PETase_RM_R | TGGTGGTGATGGTGGTGCTCGAG |
| TPA sensor_pDonor_F | GTCATTTCCGGGGATCCAGTGCCATTCCGCCTGAC |
| TPAsensor_pDonor_R | GCAGTAAGGACTCTAGAACTGAGCCTCCACCTAGC |
| pDonor_seq_F | CGACTGGAAAGCGGGCAGTG |
| pDonor_seq_R | GCCATTCAGGCTGCGCAACT |
| cr1-UF | CAACTGGTCACGCTGATCG |
| cr1-DR | CTGTTCAACCTGGTCCAGCC |
| cr2-UF | GCCATGCTAACTCGTCCAAAC |
| cr2-DR | GCGTATGAAGAGCGGTGAGC |
| cr3-UF | CCCAACTTAATCGCCTTGCA |
| cr3-DR | CTTCAGCCTCCAGTACAGCG |
| cr4-UF | CTGCATGTACGTGCCTTCC |
| cr4-DR | GCTACCGAAGGGACTATTCCTC |
| cr5-UF | CAGAGATGGAAATTACCCTGCAAG |
| cr5-DR | CGTTTGCCGTTCAACAACG |
| cr6-UF | GGTGCAAGTATCATCACCACTTC |
| cr6-DR | GTCGATTACGAGACGTTGATGAAG |
| cr7-UF | CTGGTATCGTAAAATGGTTCAACG |
| cr7-DR | CCATGTCCTGACCTGGGTTC |
| cr8-UF | TCTTCCGCATTACCCTGGTG |
| cr8-DR | TACACTGAACTCTCTGGCTCAC |
| ThermoPETase_S269T_F | ACA ACCTTTGCCTGTGAAAATCCTAACTCGA |
| ThermoPETase_S269T_R | ATAGCGGGTATCGTTATCCATGAAACGTTTCAT |
| FastPETase_S269T_F | ACAACATTTGCTTGCGAAAATCCAAACTCG |
| FastPETase_S269T_R | ATAACGCGTATCATTGTCCATAAAACGCTTCATC |

| | |
|---|---|
| HotPETase_S269T_F | ACA CAATTTGCTTGTGAAAATCCTAATTCCACTGC |
| HotPETase_S269T_R | GTAGCGGGTATCGTTGTCCATAAAACGC |
| DepoPETase_S269T_F | ATATACTACTTTCGCTTGTGAGAATCCAAATAGCA |
| DepoPETase_S269T_R | CTAGTATCATTGTCCATGAAATATTTCATCCAGGC |
| IsPETaseC273A_F | GCGGAGAATCCCAACAGCACACGC |
| IsPETaseC273A_R | GGCGAAGGTTGAGTAACGGGTGTC |
| IsPETaseC289A_F | GCGTCCCTCGAGCACCACCATCAC |
| IsPETaseC289A_R | GTTCGCGGTGCGAAAATCCGACA |
| FastPETase_C273A_F | GCGGAAAATCCAAACTCGACTGCCGTG |
| FastPETase_C273A_R | AGCAAATGTTGAATAACGCGTATCATTGTCCAT |
| FastPETas_C289A_F | GCTAGTCTCGAGCACCACCACCACCAC |
| FastPETas_C289A_R | GTTCGCGGTACGGAAATCGGACAC |

### Supplementary Note

We hypothesized that some of these effects may be the consequence of interference with the formation of the disulfide bond. The disulfide bond between Cys273 and Cys289 (DS2) is commonly thought to be conserved in both type I and type II PET degrading enzymes [7,8], but the effect on the biochemical characteristics has not been elucidated. The recent discovery of a novel PET hydrolase bbPET0069 [9], which lacks disulfide bonds, also prompts us to reconsider the exact role of this disulfide bond in the PET degrading activity. We first examined the electron density maps of the disulfide-bridges in all 12 PDB entries of the wild-type *Is*PETase. We found that in 7 out of 12 structures CYS289 displays signs of an alternative location, while only 2 in the disulfide-bridge between Cys203-Cys239, which is located close to the catalytic site (Supplementary Fig. 7). In addition, in one entry (PDB ID: 8GU5) the bridge is not formed at all, and when the electron density maps of other structures are overlaid of the atoms of 8GU5, the alternative density peaks show good correspondence with the sulfur atom of C289 in 8GU5 (Supplementary Fig. 8), indicating that in the crystals, in a substantial fraction of proteins the bridge is not present. Since sufficient mutation experiments has proved that the disruption of the other pair of disulfide bonds (C203-C239, DS1) reduced to little or no activity towards PET film [10, 7], the band shift of *Is*PETase, M2.9, ThermoPETase, Thermo_S269T, FastPETase and Fast_2.9 in Click-PEGylation [11] assay (Supplementary Fig. 6a) could most likely result from the reduction of C273-C289 disulfide bond.

Chen [12] made disulfide bonds absent double mutant (C241A/C259A) of *Thermobifida fusca* cutinase, which could also degrade PET. The catalytic efficiency of purified C241A/C259A cutinase decreased to 71.0%, and the thermostability was also reduced significantly. Similarly, we did single mutations of C273A or C289A on *Is*PETase and FastPETase, and compared the activity of the mutants with *Is*PETase and FastPETase towards PET film. Surprisingly the mutation of C273A or C289A did cause significant changes to *Is*PETase, while causing complete activity loss of FastPETase. We infer that the C273-C289 disulfide bonds are likely to play a regulatory rather than structural role in *Is*PETase and its variants, and allosteric regulation of protein activity by oxidizing/reducing a disulfide bridge was demonstrated for several proteins [13]. More studies will be needed to determine how the regulatory mechanism looks and how the residue at the 269 site might affect the regulation.
